# Comparative molecular life history of spontaneous canine and human gliomas

**DOI:** 10.1101/673822

**Authors:** Samirkumar B. Amin, Kevin J. Anderson, C. Elizabeth Boudreau, Emmanuel Martinez-Ledesma, Emre Kocakavuk, Kevin C. Johnson, Floris P. Barthel, Frederick S. Varn, Cynthia Kassab, Xiaoyang Ling, Hoon Kim, Mary Barter, Chew Yee Ngan, Margaret Chapman, Jennifer W. Koehler, Andrew D. Miller, C. Ryan Miller, Brian F. Porter, Daniel R. Rissi, Christina Mazcko, Amy K. LeBlanc, Peter J. Dickinson, Rebecca Packer, Amanda R. Taylor, John H. Rossmeisl, Amy Heimberger, Jonathan M. Levine, Roel G. W. Verhaak

**Author notes:** MedVet Medical and Cancer Center for Pets, Columbus, OH, USA. Department of Pathology, Division of Neuropathology, and O’Neal Comprehensive Cancer Center, University of Alabama at Birmingham, Birmingham, AL, USA. These authors have contributed equally (Kevin J. Anderson, C. Elizabeth Boudreau, Emmanuel Martinez-Ledesma). Co-senior authors (Amy Heimberger, Jonathan M. Levine, Roel G. W. Verhaak).

## Abstract

Sporadic gliomas in companion dogs provide a window on the interaction between tumorigenic mechanisms and host environment. We compared the molecular profiles of canine gliomas with those of human pediatric and adult gliomas to characterize evolutionarily conserved mammalian mutational processes in gliomagenesis. Employing whole genome-, exome-, transcriptome-and methylation-sequencing of 81 canine gliomas, we found alterations shared between canine and human gliomas such as the receptor tyrosine kinases, p53 and cell cycle pathways, and *IDH1* R132. Canine gliomas showed high similarity with human pediatric gliomas per robust aneuploidy, mutational rates, relative timing of mutations, and DNA methylation patterns. Our cross-species comparative genomic analysis provides unique insights into glioma etiology and the chronology of glioma-causing somatic alterations.

**Significance:** Diffuse gliomas are the most common malignant brain tumors, with high-grade tumors carrying a dismal prognosis. Preclinical models have proven themselves as poor predictors of clinical efficacy. Spontaneous glioma in dogs provides an attractive alternative model, because of their comparable tumor microenvironment and tumor life history. We determined the similarities and differences between human and canine gliomas through genomic profiling, and leveraged our datasets to identify conserved somatic drivers, mutational processes and temporal ordering of somatic glioma events across species. We show that canine gliomas resemble human gliomas at (epi-)genetic levels and are more reminiscent of pediatric than adult disease, thus rationalizing sporadic canine glioma as a preclinical model tailored to measuring treatment efficacies in patients with canine or human glioma.

## INTRODUCTION

The natural history of cancer is marked by temporal acquisition of diverse genetic and epigenetic aberrations. The inevitable intratumoral and inter-patient heterogeneity among evolving cancer cells poses a major obstacle in our understanding of cancer evolution and designing effective treatment strategies (Alizadeh et al., 2015). Recent developments in high-throughput lineage tracing, organoid cultures, and patient-derived xenografts have provided better resolution of heterogeneity and driver events. Nonetheless, in the absence of natural host response, preclinical in vitro and rodent models are unable to fully recapitulate a spontaneously evolving tumor’s life history. This limitation challenges the accuracy of predicting therapeutic responses in these preclinical models, especially response to immunotherapies (Buque and Galluzzi, 2018).

Somatic evolution of cancers may follow convergent patterns across mammalian species by selecting cells that carry beneficial mutations in highly conserved regions, i.e., genes and their regulatory noncoding regions enabling one or more cancer hallmarks (Hanahan and Weinberg, 2011). Unlike induced cancer models, comparative genomics of spontaneous tumors across species provides a unique advantage to identify defects in such shared, evolutionarily constrained regions (Lindblad-Toh et al., 2011) and to evaluate the importance of host context in the tumor’s evolution. In addition to their natural tumorigenesis, spontaneous cancers in dogs are marked by the presence of a fully functional tumor microenvironment (Khanna et al., 2006; LeBlanc et al., 2016). Cancer cells are subject to clonal selection and drift, and the resulting tumor is molded by selection pressure from the tissue context (DeGregori, 2017; Fortunato et al., 2017). This Darwinian adaptation may select for somatic alterations in evolutionarily conserved regions in both dogs and humans that are relevant to tumorigenesis.

Sporadic gliomas occur in companion dogs at frequencies similar to those in humans (Snyder et al., 2006; Song et al., 2013). Genomic characterization of canine glioma has a distinct merit, in that the age distribution at the time of diagnosis is comparable with that seen in human pediatric disease, but the animals are in the adult stage of life. This seeming conundrum in fact creates an opportunity to compare somatic drivers and their relative timing in canine glioma with those in human glioma. Studies involving comparative genomics of spontaneous canine cancers have already enabled identification of breed-specific, disease-risk loci under strong evolutionary constraints and with known roles in human cancer, e.g., germline *FGF4* retrogene expression in chondrodysplasia (Parker et al., 2009), somatic *BRAF* V600E mutation in canine invasive transitional cell carcinoma of the bladder (Decker et al., 2015b), recurrent somatic *SETD2* mutations in canine osteosarcoma (Sakthikumar et al., 2018), and *TP53* pathway alterations in canine melanoma (Hendricks et al., 2018; Wong et al., 2019). Earlier studies in canine gliomas have characterized somatic copy number alterations syntenic with those in human adult gliomas (Dickinson et al., 2016), and have identified genetic susceptibility factors near genes such as *CAMKK2*, *P2RX7*, and DVL2 (Mansour et al., 2018; Truve et al., 2016).

Here, we have performed comparative genomic, transcriptomic, and epigenetic profiling across three population structures, canine glioma, human adult and human pediatric glioma, in order to study somatic evolutionary traits of glioma across two species and in different age groups. We leveraged genomic profiles to infer molecular life history in order to understand cross-species convergent evolution of glioma (Aktipis et al., 2013; Stearns, 1992).

## RESULTS

### Human glioma driver events are frequently found in canine disease

We performed whole genome, exome, transcriptome, and methylation sequencing (370 libraries) on canine gliomas (n=81) and germline (n=57) samples from 81 dogs, with all samples obtained via necropsy. Using the recently updated criteria for diagnostic histopathological classification (Koehler et al., 2018), 42 cases were classified as oligodendroglioma, 24 cases as astrocytoma, 10 cases as glioblastoma, and 5 cases as of undefined glioma pathology (Table S1). We defined a common set of 77 cases where whole genome and exome data was available with minimum of 30X coverage in exome regions (Table S1, STAR Methods). From mutation calls derived from all 77 cases, we detected somatic mutational driver events in all 77 canine gliomas using dNdS (Martincorena et al., 2017), MuSiC2 (Dees et al., 2012), and a semi-supervised comparison with known cancer drivers in human adult and human pediatric cancers (Bailey et al., 2018; Gröbner et al., 2018; Ma et al., 2018)(Figure 1A, Table S2 and S3, STAR Methods). Mutations in genes associated with human pediatric (Mackay et al., 2017) and adult glioma (Brennan et al., 2013; Ceccarelli et al., 2016) such as the *TP53*, *PDGFRA*, *PIK3CA* and *EGFR* were observed (Figure S1A). Recurrent hotspot and mutually exclusive mutations with high oncogenic impact according to the Catalogue of Somatic Mutations in Cancer (COSMIC) database (Tate et al., 2019) were observed in *PIK3CA* H1047R/L (n=8), *PDGFRA* K385I/M (n=6), *IDH1* R132C (n=3), and *SPOP* P94R (n=1; 1 shared with *PIK3CA* H1047R) (Figure 1B). These mutations were also identified as being under positive selection or as significantly mutated genes using the dNdS (Martincorena et al., 2017) approach (Table S2) and thus indicating driver mutations of canine gliomas. Mutations affecting the *IDH1* R132 codon are a defining characteristic of low-grade adult gliomas (TCGA_Network et al., 2015) and were detected infrequently in pediatric and canine gliomas (n = 3/77). Overall, 39/77 (51%) of canine gliomas carried at least one significantly mutated gene. This proportion was similar to published findings in human pediatric tumors (57%, chi-square p-value: 0.12) (Gröbner et al., 2018) but contrasted with the frequency at which adult gliomas contain at least one significantly mutated gene alteration (93%, chi-square p-value: 1.090e-08). To demonstrate similarity between canine gliomas and human gliomas, we summarized the level of coding mutations in gene sets reflecting previously reported cancer hallmarks (Table S4). We tallied weighted pathway contributions per cohort (canine, adult, pediatric) by the number of coding mutations within each cohort and genes per pathway. Adult glioma is commonly separated into subtypes on the basis of IDH mutation as well as chromosome arm 1p and 19q deletion, resulting in three subtytpes: 1. IDH wild type (IDHwt); 2. IDH-mutant with codeletion (IDHmut-codel) and 3. IDH-mutant without codeletion (IDHmut-non-codel)(Louis et al., 2016). We have adopted these three categories for comparisons between canine glioma and human adult glioma. We found that canine and pediatric gliomas showed most concordance, with adult gliomas showing increased frequency of gene mutations in cancer hallmarks such as deregulating cellular energetics (attributed to lack of the IDH-mutant subtype), genomic instability, and resisting cell death (Figures 1C, S1B). Canine gliomas showed the most significant difference in avoiding immune destruction hallmark compared to human pediatric and adult IDHmut-non-codel cohort (Table S5). Epigenetic drivers were observed in all populations, most commonly in pediatric gliomas (72/217, 33%), followed by canine gliomas (n= 22/73, 30%), and adult *IDH1*-wild type gliomas (95/374, 25%)(Figure 1C). Telomere maintenance is the defining characteristic of human adult gliomagenesis (Barthel et al., 2018) but is not prevalent in canine (Figure S1C, S1D) or in pediatric high-grade gliomas (5/326, 1.5%)(Mackay et al., 2017). Correspondingly, neither significant changes in telomerase length nor expression changes in telomere pathway genes within canine tumors were observed (Figure S1E).

**Figure 1:**
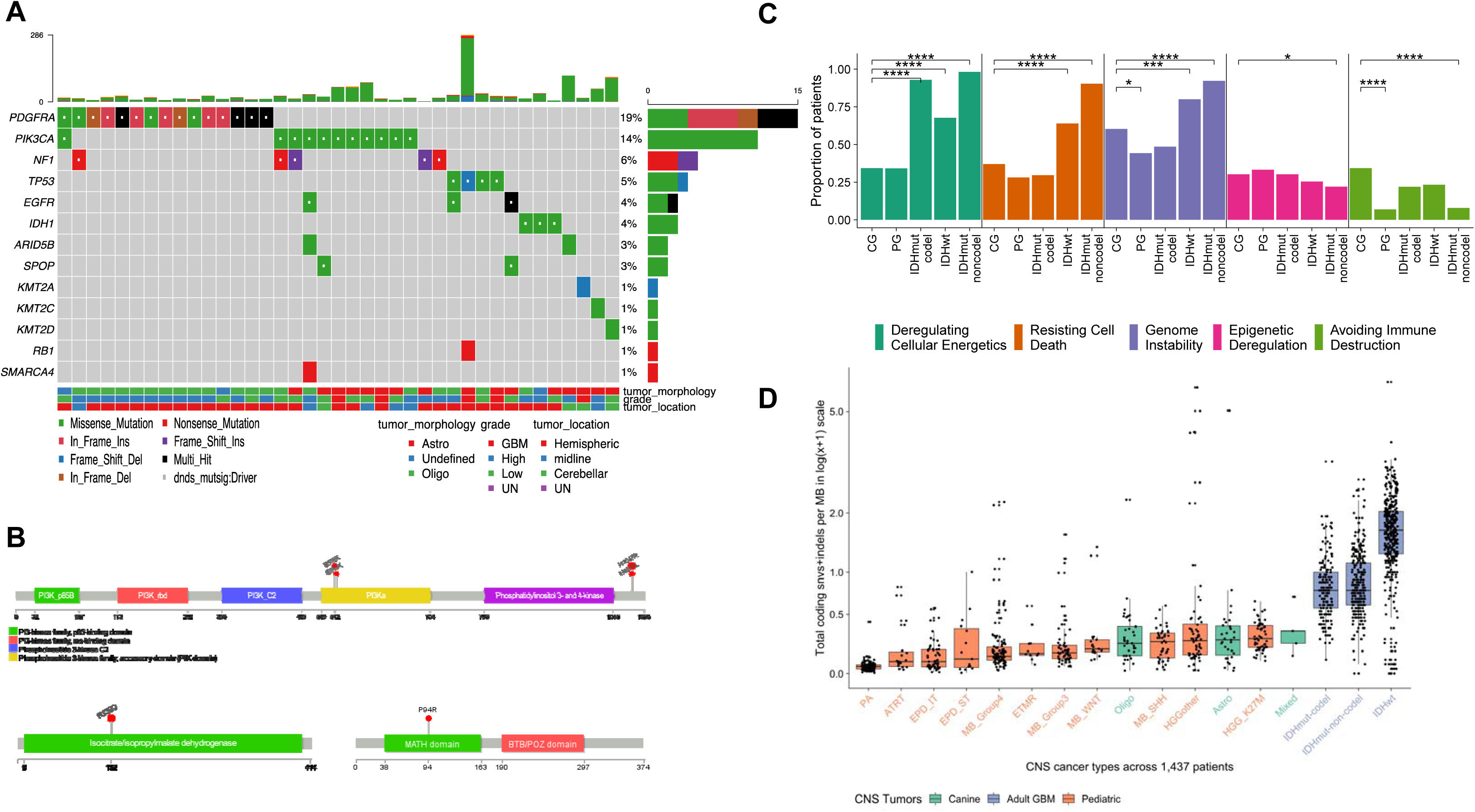
Comparative somatic landscape of canine and human gliomas. **A)** Somatic variants in canine gliomas. Top bar plot shows patient-wise frequency of somatic variants and right sided barplot shows gene-wise frequency of somatic variant types. Bottom annotations show relevant patient-specific annotations. Coding mutations of all COSMIC cancer genes is in figure S1A. UN: Undefined **B)** Gene lollipop plots showing recurrent hotspot mutations for three genes: *PIK3CA*, *IDH1*, and *SPOP*. All of hotspot mutations are validated COSMIC mutations in human cancers. **C)** Hallmark enrichment of coding mutations in cancer genes across human adult (IDHwt, IDHmut-codel, IDHmut-noncodel), pediatric (PG) and canine (CG) gliomas. Y-axis represents proportion of patients in the respective cohort harboring mutations in selected five hallmarks. Two-sided Fisher’s exact test used for comparison of proportions between cohorts. P-values less than the threshold (p < 0.05) are shown (*: < .05, **: < .01, ***: < .001, ****: < 0.0001). Hallmark plots showing all hallmarks is in figure S1B. **D)** Somatic mutation rate across canine and human brain tumors: Boxplot showing somatic mutation rates as coding mutations per megabase in log1p or log(x+1) scale. X-axis shows each of 11 different types of pediatric brain tumors (Grobner 2018), canine glioma (Oligo, Astro, Undefined), and adult gliomas separated by *IDH1/2* mutation and 1p/19q codeletion status (far right). PA: pilocytic astrocytoma; ATRT: Atypical Teratoid Rhabdoid Tumor; EPD_ST: ependymoma supratentorial; ETMR: embryonal tumors with multilayered rosettes; MB: medulloblastoma; HGG: high grade glioma. Tumors are sorted in ascending order by increasing mutation rate. Extended mutation rate plots are in figure S1F.

We compared somatic alteration rates across the canine and human pediatric and adult cohorts using coding mutation rates from 4,761 patients (Bailey et al., 2018; Ceccarelli et al., 2016; Gröbner et al., 2018; Ma et al., 2018) (STAR methods). The somatic mutation rate of canine glioma (0.25 coding mutations per megabase; 95% CI: 0.13-0.37) was similar to that of human pediatric gliomas (Figure 1D), including high-grade canine gliomas had comparable mutation rates to that of pediatric high-grade gliomas (0.26 coding mutations per megabase; Wilcoxon p-value: 0.21 or lesser; Figure S1G), but significantly lower than in human adult IDH1/2-mutant and IDH wild-type gliomas (0.66 and 1.89 coding mutations per megabase, respectively; Wilcoxon p-values of 2.2E-16 or lesser). Low mutation burden has been linked to fewer mutations in cancer-driving genes (Martincorena et al., 2017) and may explain the relative paucity of significantly mutated genes observed in canine gliomas, including weaker positive selection (q > 0.1) for known and mutated cancer genes (n=78; Figure S1H). These results demonstrate that the landscape of somatic single-nucleotide variants is similar to human glioma, and that canine glioma aligns more closely with human pediatric glioma than with adult disease.

### Aneuploidy is a major driver of canine and pediatric high-grade glioma

We compared the DNA copy number landscape of glioma across species with a focus on the >50% of canine gliomas without evidence of significantly mutated genes. No focal copy number amplifications were detected among canine gliomas. Human glioma tumor suppressor gene *CDKN2A/B* was homozygously deleted in 7/56 (13%, 6/7 astrocytomas), and *PTEN* in 2/56 (4%) of canine glioma genomes (Figures 2A, S2A). Together, 54/77 (70%) patients with canine glioma contained somatic mutations and focal copy alterations in known human glioma drivers. Contrasting with the limited presence of focal DNA copy number alterations was the high frequency of arm-level copy gains (canine chromosomes 7q, 13q, 16q, 20q, 34q, 35q, and 38q) and arm-level losses (canine chromosomes 1q, 5q, 12q, 22q, and 26q) (Figure S2B). The most frequent arm-level alteration comprised the shared syntenic regions of glioma drivers *PDGFRA*, *KIT*, *MYC* (Figure S2C) and typically resulted in more than four copies of these genes (canine 13q+; 19/77 cases, 25%). Other common arm-level alterations included *PIK3CA* (canine 34q+) and the *HIST1* cluster (canine 35q+) as well as hemizygous loss of heterozygosity of tumor suppressor genes, *TP53*, *RB1*, and *PTEN* (Figure 2A).

**Figure 2:**
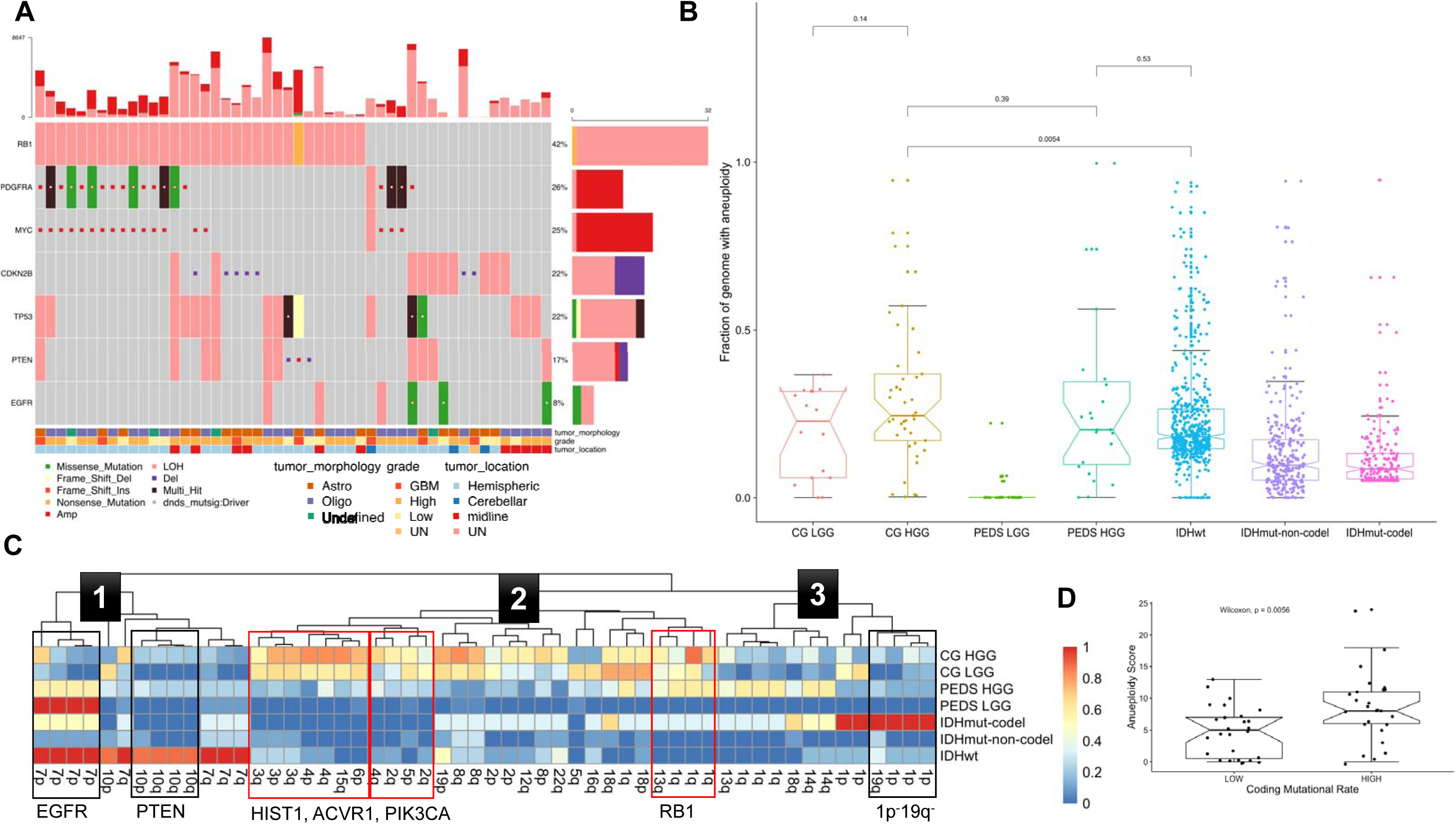
Aneuploidy is a major driver of high-grade gliomas. **A)** Focal somatic copy alterations in canine gliomas (n = 57). Squared symbol in cell suggests either amplification (>4 copies) or deep deletion based on GISTIC2 gene-level table (Methods). Top bar plot shows patient-wise frequency of somatic variants and copy number alterations and right sided barplot shows driver-wise frequency of somatic variant types, including copy number alterations. Bottom annotations show relevant patient-specific annotations. UN: Undefined **B)** Comparative aneuploidy score: Boxplots showing fraction of genome with aneuploidy (Y-axis) for human adult glioma, separated by IDH-mutation and 1p/19q codeletion status; pediatric (PEDS), and canine (CG) cohorts, further divided as low grade (LGG) or high grade (HGG). Pairwise p-value are calculated using Wilcoxon non-parametric test. **C)** Aneuploidy metrics across shared syntenic regions of canine and human genome: Heatmap showing comparative aneuploidy across three cohorts. Each column shows the aneuploidy fraction (% of aneuploid genome) for a given shared syntenic region with **label corresponding to human chromosome arm**. Each of seven rows represent subgroups of canine and human gliomas based on histopathology (for canine and pediatric) and IDH-mutation and 1p/19q codeletion status. Numbered boxes on the dendrogram represents three aneuploidy cluster described in the main text. **D)** Boxplot comparing coding mutation rate to aneuploidy score for canine gliomas: Pairwise p-value are calculated using Wilcoxon non-parametric test.

We quantified prevalence of aneuploidy across the canine, human pediatric and adult glioma populations (Taylor et al., 2018). For copy-number estimation, matched tumor-normal whole genome sequencing profiles from canine (n=57) and pediatric gliomas (n=50)(Ma et al., 2018), and Affymetrix SNP6 profiles for adult gliomas (n=969)(Ceccarelli et al., 2016) were analyzed (STAR Methods). Aneuploidy metric was based on the proportion of copy-number segmented genome, which is non-diploid (STAR Methods). Canine gliomas, independent of tumor grade, and pediatric high-grade gliomas showed comparable aneuploidy (25% and 20% of genome aneuploidy, respectively), which was significantly higher than that in human low-grade pediatric glioma (near-euploid genome) and adult glioma (9-18% of genome aneuploid) (Figure 2B). We then searched for aneuploidy within syntenic regions, which may be subject to selection pressure during gliomagenesis. We mapped canine chromosome arms to their human counterparts and used unsupervised hierarchical clustering of the most variable syntenic aneuploid regions to identify regions of shared aneuploidy (Figure 2C). The analysis revealed three aneuploidy clusters. The first cluster consisted of arm-level aneuploidy of human 7p (*EGFR*) and 10q (*PTEN*) arms characteristic of human adult *IDH1* wild-type (100%, 89% respectively) and pediatric high-grade gliomas (57%, 23%) for which the canine, pediatric high-grade, and pediatric low-grade gliomas showed 10%, 18% and 0% alterations, respectively. None of three *IDH1* mutant canine gliomas shared these syntenic aberrations, suggesting a mutually exclusive pattern as observed in human gliomas. The second cluster consisted of human 2q/6p, which contains the *ACVR1* and the *HIST1* genes that are frequently mutated in pediatric high-grade gliomas, with significant enrichment within H3.1K27M diffuse intrinsic pontine glioma, along with mutations in RTK pathway genes (*PIK3CA*, *PIK3R1*)(Mackay et al., 2017), which showed loss of the syntenic human 2q/canine 36q region (containing *ACVR1*) within 44%, 23% and 23% of canine high-grade, canine low-grade, and pediatric high-grade gliomas, respectively. In contrast, this alteration was not observed in human pediatric or adult IDH mutant glioma but was present in 12% of IDH wild-type adult gliomas. Similarly, human chromosome arm 6p/canine chromosome arm 35q, containing the *HIST1* gene cluster, was frequently amplified in canine high-grade glioma (75%) and low-grade glioma (67%) and pediatric high-grade gliomas (29 %) but not in pediatric low-grade or adult gliomas. The third cluster consisted of human 1p/19q codeletions seen commonly in adult *IDH1* mutant gliomas but observed in 15-30% of cases in canine respectively human pediatric cohorts. Comparing the aneuploidy score among canine gliomas with high vs. low coding mutational rate showed significant (Figure 2D; Wilcoxon p-value: 0.006) increases in aneuploidy among patients with a high mutational rate, suggesting that an underlying mutational process promotes genomic instability during gliomagenesis.

### DNA damage-related mutational processes shape somatic driver landscape and maintain genomic instability

We leveraged known mutational signatures from adult (COSMIC v2, 1 to 30) and pediatric cancers (T1 to T12) to estimate and compare underlying mutational processes across canine and human gliomas (Alexandrov et al., 2013; Gröbner et al., 2018; Ma et al., 2018). The most enriched signatures across all canine gliomas (Figure S3A) were associated with aging (COSMIC signature 1, pediatric signature T1), mismatch repair deficiency (COSMIC signature 15), APOBEC-AID (COSMIC signature 2, 9), homologous repair defect signatures (COSMIC signature 8, pediatric signature T3), and signatures with unknown relevance, COSMIC signature 12, T10, and T11. Among the nine canine gliomas with the highest mutation rates (median coding mutation rate of 0.55 per megabase)(Figure 3A), there was significant (Wilcoxon p-value: 0.025) enrichment of two additional mismatch repair signatures (T9 or COSMIC signature 3, 6)(Figure S3B). A frameshift indel in mismatch repair gene *MSH6* was detected in one case with an outlier mutation frequency (coding mutation rate of 5.04 per MB)(Figure S3C). Among the remaining cases (median coding mutation rate of 0.25 per MB), homologous repair defect or ‘BRCAness’ signatures (COSMIC signature 3 or pediatric signature T3, COSMIC signature 8 or pediatric signature T6) were the second most prominent signatures after clock-like signatures (COSMIC signature 1, 5). Homologous repair defect signatures have been reported to be enriched in pediatric high-grade gliomas with higher genomic instability (Gröbner et al., 2018). The known human signatures were validated by clustering *de-novo* constructed signatures for all three cohorts (canine, human adult, and pediatric gliomas). Independent of cohort type, we observed significant cosine similarity (more than 0.8; Figure 3B, Figure S3D) of *de-novo* signatures with known homologous repair defect mutational processes (COSMIC signature 3/pediatric signature T3, COSMIC signature 8/pediatric signature T6 among others), further implying a role for these mutational processes in cross-species gliomagenesis.

**Figure 3:**
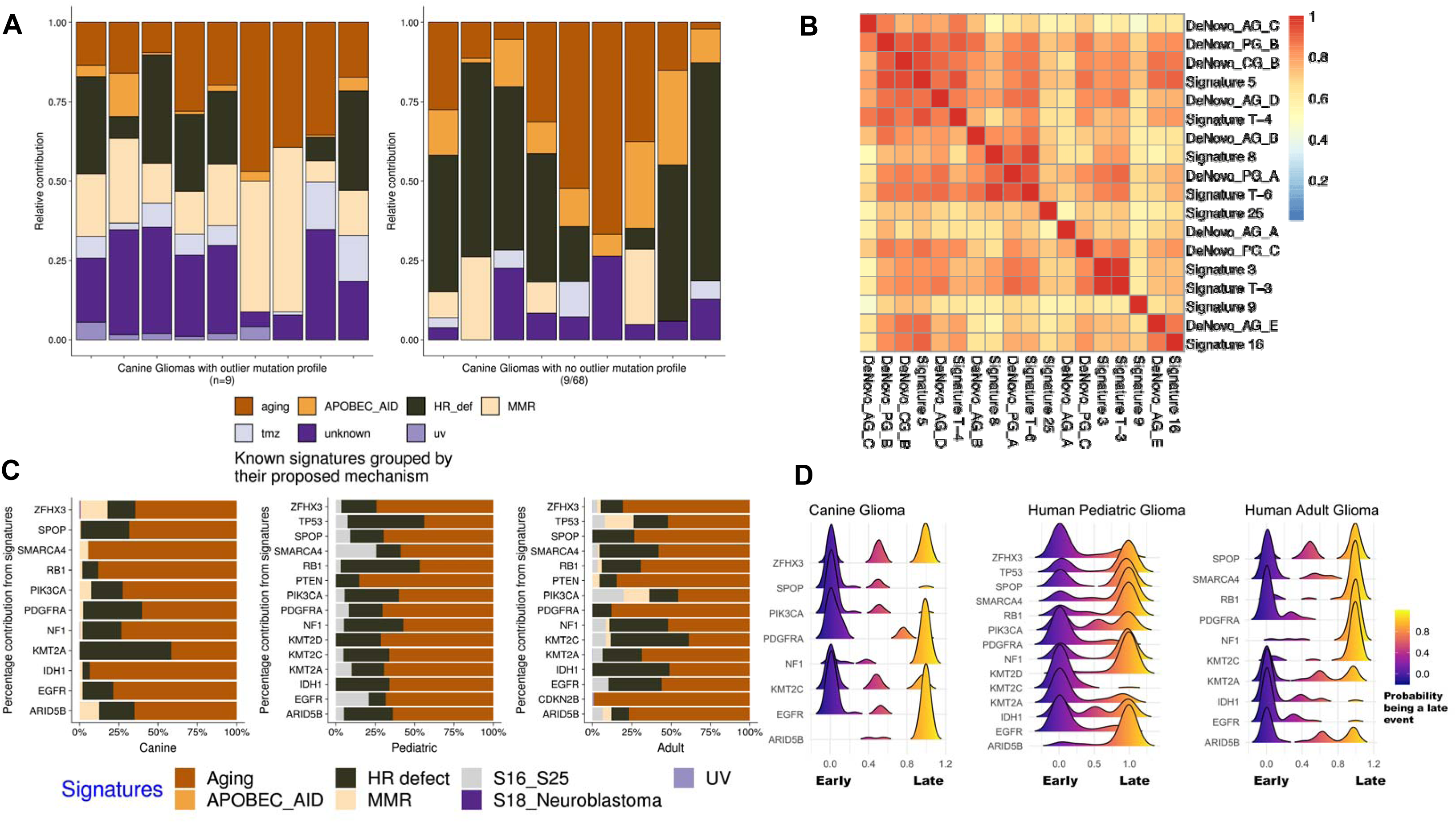
Molecular life history analysis using mutational signatures and timing analysis. **A)** Deconvolution of known human mutational signatures on canine glioma somatic variant data: Stacked barplots shows relative contribution of known human mutational signatures in individual canine patients. Signature contributions were aggregated based on their grouping into proposed mechanism. Only top signatures with relative contribution more than third quartile per sample is shown in the plot. Plot on the left side shows 9 cases with highest mutational frequency (based on outlier mutational profile, STAR methods) and plot on the right side shows 9 representative cases with median signature contribution within inter-quartile range. Table S6 provides mapping between signature and proposed mechanisms. Signatures with no proposed mechanism are grouped into the *unknown* category. **B)** Hierarchical clustering of cosine similarities between known human mutational signatures and de-novo signatures constructed using available whole genome data from canine (CG), pediatric (PG), and adult (AG) data. Higher cosine similarity (red color) indicates higher resemblance of de-novo signature to known mutational signature. Only one of three cluster group is shown here and the complete clustering is in figure S3D. **C)** Horizontal stacked barplots represent percentage contribution of signature groups (X-axis) for somatic driver mutations (Y-axis) found in canine and human gliomas. Each of seven signature groups are combination of one of more known human signatures (Table S6). APOBEC_AID: activation-induced cytidine deaminases; HR defect: homologous repair defect; MMR: mismatch repair; S16_S25 and S18_Neuroblastoma: signatures previously described by Gröbner et al. **D)** Molecular timing of somatic drivers across canine and human gliomas: Stacked density plots, one per each of three cohorts, shows probability (X-axis) of a driver event (Y-axis) being a late event in tumor evolution and value of <0.5 being an earlier event. Density plots for each driver event were calculated based on pairwise winning probability (where win is defined a early event) as used in sports statistics (Bradley-Terry model). Winning probabilities were subtracted from 1 to display early events on the left side of the plot.

Next, we determined the relative contribution of mutational processes (with deconvoluted signatures as a proxy) in generating mutations within significantly mutated genes to identify the dominant mutational process(es) active during tumor evolution (Figure 3C). Although clock-like processes (COSMIC signature 1, 5) largely contributed to an age-related increase in mutations, including in driver genes, we found that homologous repair defect signatures (COSMIC signature 3, 8) contributed (27%, 21/77 cases) to driver mutations across all three cohorts, emphasizing that homologous repair defect can not only serve as a potential source for driver mutations but also fuel progressive genomic instability along with observed high aneuploidy (Blank et al., 2015; Targa and Rancati, 2018) in high-grade gliomas across all three cohorts.

### Comparative molecular timing analysis highlights context-specific early and late drivers of gliomagenesis

We inferred the sequential order of somatic alterations during gliomagenesis by estimating clonality of glioma driver events (Figure 3D) (Jolly and Van Loo, 2018; Shinde et al., 2018). In brief, significantly mutated genes were timed as occurring early (clonal) to late (subclonal) during tumor evolution based on their cancer cell fraction after accounting for tumor purity, ploidy, and copy number status (Methods). We observed clonal *PDGFRA* and *EGFR* mutations as the only shared and early event across all three cohorts. Subsequent whole chromosome 13 amplification bearing the *PDGFRA* mutant allele marked the emergence of the most recent common ancestor (MRCA) in six canine gliomas (Figure S3E), which grew to be a dominant clone at the time of diagnosis. *IDH1* mutation marks an initiating event in IDH1-mutant human gliomas (Barthel et al., 2018). Correspondingly, *IDH1* mutations were ubiquitously timed as an initiating event (CCF > 0.9) in three canine and three human adult *IDH1* mutant cases, and as an early event in one case of pediatric glioma (CCF = 0.83). We observed *NF1* frame-shift mutations mostly as a late event across all cohorts whereas *PIK3CA* mutations appeared as an early event for canine and human pediatric gliomas. Although the relatively uniform timing patterns of these known glioma drivers suggest convergent evolution in varied contexts, i.e., presence of hotspot mutations in shared drivers (*PDGFRA*, *PIK3CA*) during clonal evolution of glioma across two species and different age-groups, we also observed an oscillating pattern of timing and consequent underlying natural selection for a set of epigenetic drivers in the lysine methyltransferase (MLL) family (Rao and Dou, 2015). *MLL3* (*KMT2C*) gene mutations were clonal events in canine and pediatric gliomas but subclonal in adult gliomas, whereas *ARID5B* mutations showed the inverse pattern (Figure 3D). MLL family genes include some of the most commonly mutated genes in pediatric cancers, including gliomas (Huether et al., 2014; Sturm et al., 2014) but not in adult gliomas (Bailey et al., 2018).

### Canine gliomas are classified as pediatric glioma by DNA methylation

We hypothesized that epigenetic deregulation in canine gliomas may carry a tumor-specific methylation pattern reflecting underlying tumor pathology, as has been observed across human brain tumors (Capper et al., 2018). We leveraged reduced representation bisulfite sequencing of canine gliomas to generate genome-wide DNA methylation profiles in order to classify canine gliomas according to a classification model widely used for human brain tumors (Capper et al., 2018). As the human brain tumor classifier was developed using the Illumina human 450k array platform, we developed a logistic regression model to enable classification of the sequencing-based canine DNA methylation profiles. We found that the model classified 35/45 (78%) of canine samples as pediatric glioma (Figure 4). Six of 45 (13%) samples were classified as IDH wild-type adult glioma, and 4/45 (9%) samples were classified as IDH-mutant adult glioma. Of the three samples carrying an *IDH1* R132 mutation, one was classified as IDH-mutant adult glioma, with a classification probability of 99%, while a second IDH-mutant sample had a relatively high classification probability for IDH-mutant adult glioma (40%), in parallel with a 57% pediatric glioma classification probability. The third sample had a low classification probability for IDH-mutant adult glioma (13%) and was classified as pediatric glioma with an 84% probability. Although the majority of canine samples were classified as pediatric glioma, no samples in the dataset were below the age of sexual maturity in canines, which is reached between 10 months and two years of age (Thompson et al., 2017). Adult human high-grade glioma tends to be restricted to the cerebral hemispheres, whereas pediatric high-grade gliomas occur throughout the central nervous system with about half of pediatric high-grade gliomas occurring in midline locations (Mackay et al., 2017). Of ten midline canine tumors (six cerebellar, four midline), eight were classified by DNA methylation as pediatric glioma and two cases were labeled as adult IDH1-mutant.

**Figure 4:**
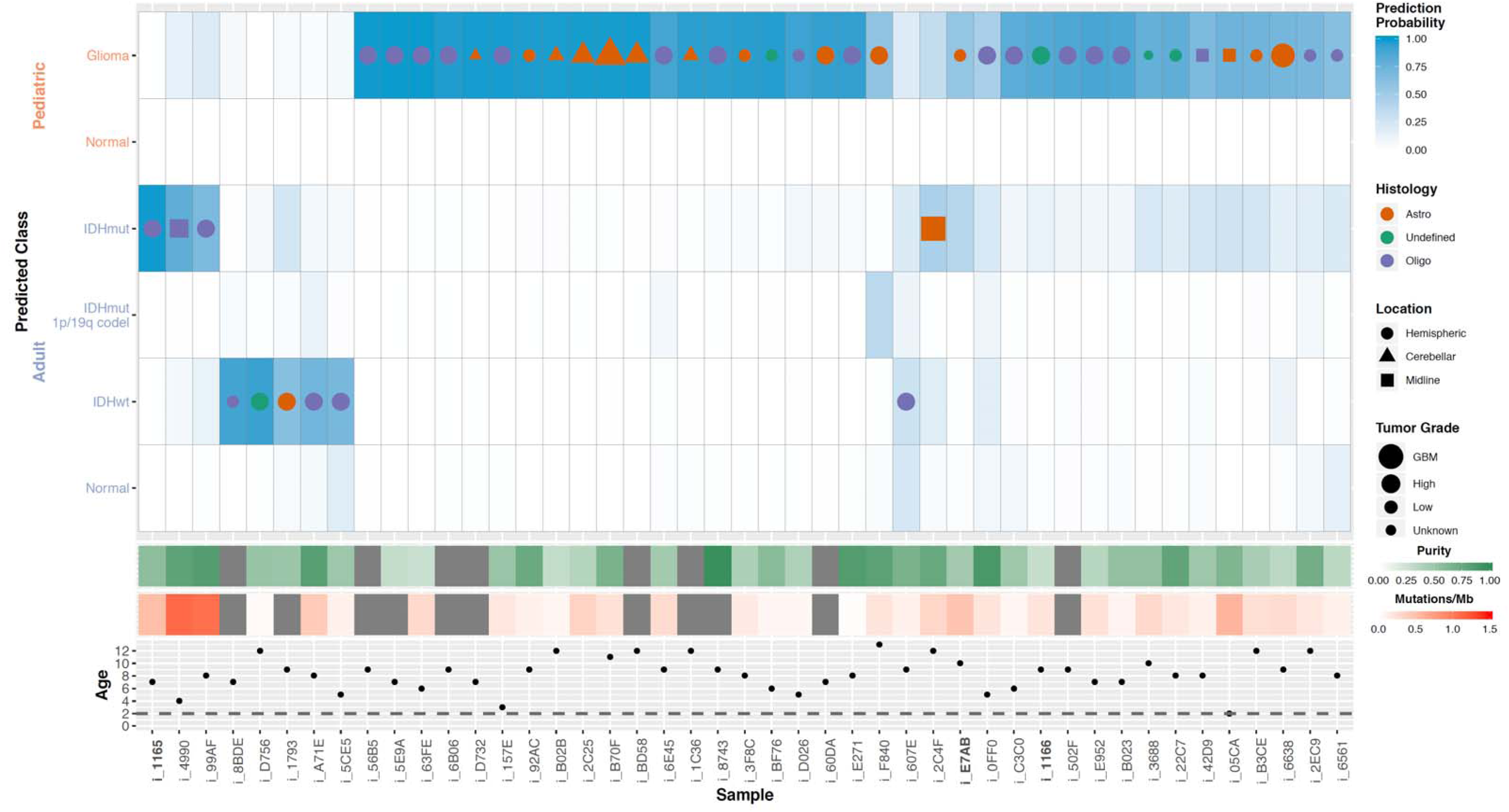
Classification of canine gliomas using human brain tumor methylation classifier. Heatmap displaying results of L2-regularized, logistic regression classification of canine methylation profiles. Each column of the heatmap represents sample, and each row in the top panel is the probability that that sample falls under a given subtype classification. The classification with the highest probability in a given sample has a symbol with symbol color, size, and shape denoting sample histology, tumor grade, and anatomical location, respectively. Panels below the probability heatmap show the tumor purity, and somatic mutation rate, and age for the samples. The horizontal line on the age subpanel denotes the age of maturity for canines (2 years).

DNA methylation profiles have been used to estimate molecular age (Pai et al., 2011). We used this approach to compare the level of age acceleration in canine and human glioma. No significant difference was observed in inferred DNA methylation age between canine tumors classified as adult glioma versus those classified as pediatric among tumors with a classification probability greater than 50% (5.945 vs 5.958, p-value 0.9125), consistent with the lack of correlation observed between canine methylation classification and chronological age. The normalized mean age acceleration was significantly higher for human pediatric glioma samples (2.5) than either human adult glioma (0.8) or canine glioma samples (−0.18) (Figure S4). Unlike human samples, the DNA methylation-inferred age did not correlate with chronological age for canine samples (Pearson correlation coefficient = 0.21), which may reflect limitations in the aging clock model derived for canids, rather than biological differences in canine tumor methylation. The DNA methylation profile of canine glioma further corroborates the evidence that glioma in dogs is generally more similar to human pediatric glioma than human adult glioma.

### Immune microenvironment

As spontaneous tumors arising in immune-competent hosts, canine gliomas represent an excellent resource through which to improve our understanding of how the immune system responds to and affects brain tumor development. To obtain a baseline understanding of how the canine glioma microenvironment compares with that of adult and pediatric gliomas, we inferred the relative immune cell fractions from human adult (n=703), pediatric (n=92), and canine glioma (n=40) RNAseq derived gene expression profiles by using the leukocyte gene signature based CIBERSORT deconvolution method (Newman et al., 2015) (Figure 5). Notably, there are many key shared immunological features between the human and canine gliomas such as the relative scarcity of CD8 T cell markers (albeit higher in high-grade gliomas), a predominance of monocyte and macrophage infiltration markers, and similar degrees of CD4 T cell, NK, and myeloid-derived suppressor cell markers, indicating that dogs with spontaneously arising gliomas may be excellent models for the testing of immune therapeutics although there may be increased toxicities associated with histamine reactions to some agents in canines. Canine gliomas exhibited two fundamental immunological differences relative to the human glioma counterpart by having significant expression of B cells and eosinophil markers (q-value < 0.05). B-lymphocyte derived plasma cells were also common among canine gliomas as well human adult gliomas, but notably absent in human pediatric tumors. The relative immune cell fractions found in each glioma type were well correlated with one another, with the low-grade pediatric glioma exhibiting the lowest correlation with high-grade canine glioma (Rho = 0.83).

**Figure 5:**
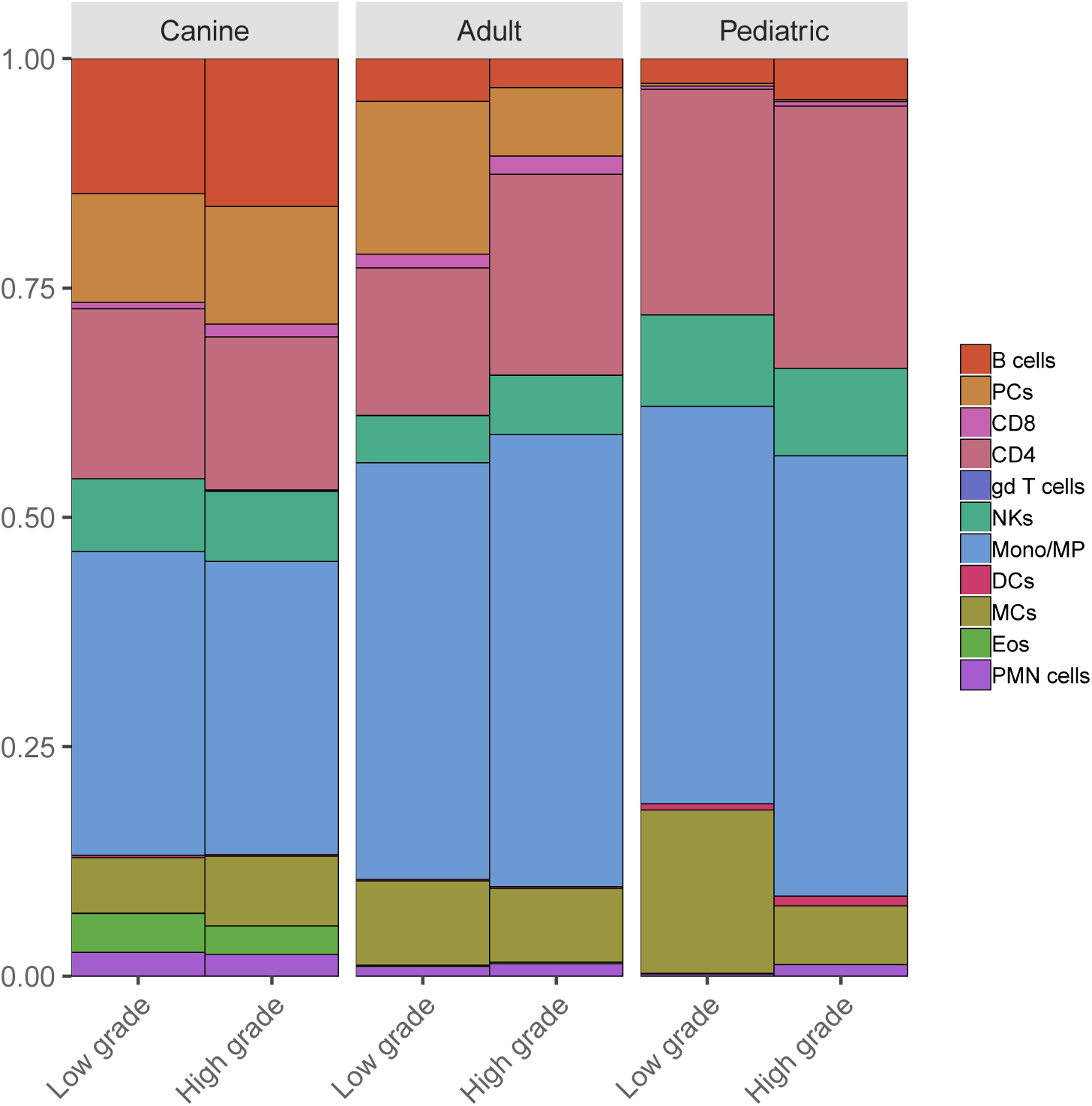
Gene expression based deconvolution of canine and human gliomas to infer immune microenvironment. Relative enrichment of estimated immune cells based on gene expression data – CIBERSORT based deconvolution of immune cells proportion in canine and human pediatric and adult gliomas. Signatures are based on human derived immune cell signature.

## DISCUSSION

Comparative genomic oncology is a robust approach for identifying evolutionarily conserved drivers and for studying the natural history of spontaneous tumors in an immune competent host, e.g., in domestic dogs (Decker et al., 2015a; Frampton et al., 2018; Tollis et al., 2017). Our cross-species analysis using comprehensive molecular profiling of sporadic gliomas highlights two key findings. First, convergent evolution of gliomas dominates between canine, human pediatric and adult gliomas, with shared molecular traits such as shared hotspot and mutually exclusive mutations in *PDGFRA* and *PIK3CA*, and in genes associated with for example the p53 and cell cycle pathways. This is further supported by aneuploidy being prevalent among canine and human pediatric high-grade gliomas, which are potentially under selection pressure within shared syntenic regions of the genome. Also, DNA damage-related mutational processes such as homologous recombination defects constitute a major source for progressive genomic instability, and generate somatic variations upon which natural selection acts to produce shared molecular and histopathological features of glioma. Second, the molecular landscape of canine gliomas resembles that of human gliomas based on the observed pattern of somatic alterations among non-shared drivers and DNA methylation patterns. Convergent evolution can reflect a footprint of adaptation to similar selective pressures (Fortunato et al., 2017). While such convergence is well-appreciated in human cancers, and in particular treatment-resistant cancers (Venkatesan et al., 2017), our observation of such molecular and phenotypic convergence across two species provides a strong indicator of variations under selective pressures exerted by the tissue or ecological context (DeGregori, 2017; Schneider et al., 2017). We note that convergent evolution should not discount a possibility of drivers unique to canine gliomas, especially within the context of germline variants (Mansour et al., 2018; Truve et al., 2016) and noncoding regulatory regions (Lindblad-Toh et al., 2011; Villar et al., 2015). Characterization of such species-specific drivers can be of much value to identify evolutionary lynchpins, which if abrogated can drive oncogenesis with similar histopathological and clinical traits. Further studies are needed to help understand how the time point at which tissue sample used in our comparative analysis were obtained, necropsy for canine samples and at diagnosis for human samples, impacts our results.

The molecular life history of a tumor is marked by multiple, often successive aberrations in genes (Armitage and Doll, 1954; Nowell, 1976). Accordingly, cancer is largely a disease of old age except in cases with early exposures to mutagens, i.e., germline or acquired defects in one or more hallmarks of cancer (Hanahan and Weinberg, 2011). The median age of occurrence for canine gliomas in our cohort was 9 years, i.e., dogs in their adult stage of life. However, we demonstrate that canine gliomas have a significantly lower somatic mutation rate, and consequently, a lower number of significantly mutated genes than adult human gliomas. Canine gliomas harbor significantly higher aneuploidy than adult human high-grade gliomas, which is more similar to human pediatric high-grade gliomas (Gröbner et al., 2018; Mackay et al., 2017). We find additional support for aneuploidy as a major driver in canine and pediatric high-grade gliomas in the observation of aneuploidy in regions of shared synteny containing the *HIST1* and *ACVR1* genes, known pediatric glioma drivers (Gröbner et al., 2018; Mackay et al., 2017), and in noting shared homologous repair defects as a mutational process that could drive genomic instability (Blank et al., 2015; Targa and Rancati, 2018). Recent efforts to engineer aneuploidy have provided better understanding on the functional role of aneuploidy and how it can be targeted in cancer (Bakhoum and Cantley, 2018; Taylor et al., 2018). Canine high-grade gliomas carrying aneuploidy, especially among syntenic regions carrying the *HIST1* and *ACVR1* genes, can be utilized as a preclinical model for such functional screening as well as to validate recent studies showing its role in immune evasion (Bakhoum et al., 2018; Davoli et al., 2017).

Tissue context and tumor microenvironment are critical factors for tumorigenesis, and current models are unable to accurately represent the development of spontaneous tumors. This renders preclinical evaluation ineffective and increases costs of clinical trials and results in minimal yields for patients. Preclinical trials of dog glioma patients enable identification of evolutionarily constrained and potentially targetable drivers, but they simultaneously benefit dogs with glioma by offering treatment options that otherwise are prohibitive due to associated healthcare costs (LeBlanc et al., 2016). Future efforts leveraging results from the comparative genomics of glioma to study immune-mediated host responses can shed light on the complex interplay between the tumor and host immune response and also aid in optimizing ongoing parallel canine clinical trials (Addissie and Klingemann, 2018) in order to improve an otherwise limited response to immunotherapies in canine and human gliomas. With respect to the immune microenvironment, differences in immune cell gene expression patterns between species could confound immune cell comparisons by under- or overestimating the presence of specific immune cell types. Despite these potential differences, comparative transcriptomic analyses of mouse and human immune cells have shown that the cells in each species exhibit a high degree of global conservation with one another, and signatures derived from murine immune cells have provided accurate immune infiltration estimates in human cancer types (Shay et al., 2013; Varn et al., 2017). Thus, the estimates in this study provide a baseline for how the relative fractions of major immune cells compare among adult, pediatric, and canine gliomas. Moving forward, signatures derived from canine immune cells will be of value in examining the presence of more specific immune cell types.

In summary, our study shows that the comparative molecular life history of gliomas details conserved drivers of glioma at both the genetic and epigenetic levels, with aneuploidy as a major hallmark of high-grade disease. Our results effectively position preclinical models of spontaneous canine glioma for use in understanding glioma drivers, and evaluation of novel therapies targeting aneuploidy, immunotherapies, with relevance to all human gliomas and pediatric disease in particular.

## ACKNOWLEDGEMENTS

This work is supported by grants from the National Institutes of Health: Cancer Center Support grants P30CA16672 and P30CA034196 and Cancer Center Support Grant Supplement 3P30CA016672-41S7 (A.B.H); R01 CA190121 (R.G.W.V.), R01 CA120813 (A.B.H); P0 1CA207206 (J.H.R.); an unrestricted grant from Agilent Technologies (R.G.W.V.); and philanthropic support from Mr. Herb Simmons (A.B.H.). E.K. is recipient of an MD-Fellowship by the Boehringer Ingelheim Fonds and is supported by the German Academic Scholarship Foundation. F.B. is supported by the JAX Scholar program and K99 CA226387; K.C.J. is the recipient of an American Cancer Society Fellowship (130984-PF-17-141-01-DMC). We thank Margaret Chapman for input on breed and clade classification, collating patient sample metadata. This work was partially supported (AKL, CM) by the Intramural Program of the National Cancer Institute, NIH (Z01-BC006161). The content of this publication does not necessarily reflect the views or policies of the Department of Health and Human Services, nor does mention of trade names, commercial products, or organizations imply endorsement by the U.S. Government.

## AUTHOR CONTRIBUTIONS

R.G.W.V., J.L. and A.B.H conceived, supervised and financially supported the study. B.B., P.V.D., J.W.K., J.L., R.P., J.H.R., and A.R.T. provided canine patient samples. J.W.K., A.D.M., C.R.M., B.P., D.R.R., C.M., and A.K.L. provided consensus histopathological classification on canine gliomas. Sample processing, quality control, and sequencing was performed by C.Y.N. and M.B.. S.B.A. and R.G.W.V. designed analysis themes. S.B.A., K.C.A., B.B., and M.C. collected patient samples and curated metadata. Data analysis was led by S.B.A. in collaboration with K.C.A., K.J.A, F.P.B., E.K., H.K., E.M.L., and F.V.. All authors participated in the discussion of the results. S.B.A. and R.G.W.V. wrote the manuscript. All co-authors discussed the results and commented on the manuscript and Supplementary Material.

## EXPERIMENTAL MODELS AND SUBJECT DETAILS

### Patient and tissue samples

Tissue samples from canine patients with gliomas were acquired with material transfer agreements from Auburn University College of Veterinary Medicine, Colorado State University, Texas A&M College of Veterinary Medicine & Biomedical Sciences, UC Davis School of Veterinary Medicine and Virginia-MD College of Veterinary Medicine. Tissue samples from resected tumor (n=77) and matched normal tissue (n=57 or paired cases) were collected at the surgical treatment or immediately following euthanasia. There were also four additional dog patients where we had adequate DNA and RNA for methylation (n=45) and RNA-seq (n=40) profiling but unable to do WGS/Exome sequencing because of failed library preparation (see Table S1). Matched normal tissue were from post-necropsy sample of contra-lateral healthy brain tissue (n=32), white blood cells (n=11), and remaining 14 samples from other tissues. Samples were archived in snap-frozen (n=33/57 paired cases; n=7/20 tumor-only cases) and Formalin-Fixed Paraffin-Embedded (FFPE, n=24/57 paired cases; n=13/20 tumor-only cases) state. Samples were then shipped to sequencing core facilities for sample preparation, quality control and sequencing (see Methods below).

### METHODS DETAILS

#### Published data sources

For comparison to human glioma, we downloaded both - raw sequencing data and processed tables for human pediatric and adult gliomas with appropriate controlled-data access agreements where needed. We used published mutation rates (Figure 1D) and mutational signatures (Figure 3) from pan-cancer datasets from adults (n=3,281) and pediatric (n=961) cohorts (Alexandrov et al., 2013; Bailey et al., 2018; Gröbner et al., 2018). For aneuploidy and molecular life history analysis (details below), we downloaded raw sequencing data and analyzed whole genomes from 53 pediatric gliomas (Ma et al., 2018; St. Jude Cloud Pediatric Cancer Genome Project, https://pecan.stjude.cloud), SNP6 data from adult gliomas – IDHwt (n=517), IDHmut-codel (n=171), and IDHmut-noncodel (n=281) cases (Ceccarelli et al., 2016), as well as whole genomes from 23 adult GBMs (Brennan et al., 2013). For coding mutation rate calculation, we used a subset of TCGA glioma set where exome/whole genome based variant calls were available: IDHwt (n=371), IDHmut-non-codel (n=268), and IDHmut-codel (n=169).

#### Sample preparation, Quality controls (QC) and Sequencing strategies

<u>DNA/RNA extraction</u> - Genomic DNA and total RNA of fresh frozen tissue and FFPE tissue from paraffin scrolls were were extracted simultaneously using AllPrep DNA/RNA Mini Kit (Qiagen) and AllPrep DNA/RNA FFPE Kit (Qiagen) according to the manufacturer’s instructions, respectively. Additional DNase treatment was performed on-column for RNA purification. <u>WGS sample preparation</u> - 200-400ng of DNA was sheared to 400bp using a LE220 focused-ultrasonicator (Covaris) and size selected using Ampure XP beads (Beckman Coulter). The fragments were treated with end-repair, A-tailing, and ligation of Illumina compatible adapters (Integrated DNA Technologies) using the KAPA Hyper Prep Kit (Illumina) (KAPA Biosystems/ Roche). For FFPE samples, 5 to 10 cycles of PCR amplification were performed. Quantification of libraries were performed using real-time qPCR (Thermo Fisher). Libraries were sequenced paired end reads of 151bp on Illumina Hiseq X-Ten (Novogene). <u>WES sample preparation</u> - Sample were prepared as described above in the WGS sample preparation, targeting 200bp with PCR amplification. Target capture was performed using SeqCap EZ Canine Exome Custom Design (canine 140702_canFam3_exomeplus_BB_EZ_HX1 probe set) (Broeckx et al., 2015) (Roche Nimblegen). Briefly, WGS libraries were hybridized with capture probes using Nimblegen SepCap EZ Kit (Roche Nimblegen) according to manufacturer’s instruction. Captured fragments were PCR amplified and purified using Ampure XP beads. Quantification of libraries were performed using real-time qPCR (Thermo Fisher). Libraries were sequenced paired end of 76bp on Hiseq4000 (Illumina). <u>RNA-seq sample preparation</u> - RNA-seq libraries were prepared with KAPA Stranded mRNA-Seq kit (Kapa Biosystem/ Roche) according to manufacturer’s instruction. First, poly A RNA was isolated from 300ng total RNA using oligo-dT magnetic beads. Purified RNA was then fragmented at 85°C for 6 mins, targeting fragments range 250-300bp. Fragmented RNA is reverse-transcribed with an incubation of 25°C for 10mins, 42°C for 15 mins and an inactivation step at 70°C for 15mins. This was followed by second strand synthesis at 16°C, 60 mins. Double stranded cDNA (dscDNA) fragments were purified using Ampure XP beads (Beckman). The dscDNA were then A-tailed, and ligated with illumina compatible adaptors (IDT). Adaptor-ligated DNA was purified using Ampure XP beads. This is followed by 10 cycles of PCR amplification. The final library was cleaned up using AMpure XP beads. Quantification of libraries were performed using real-time qPCR (Thermo Fisher). Sequencing was performed on Hiseq4000 (Illumina) generating paired end reads of 75bp. <u>Reduced Representation Bisulfite Sequencing (RRBS) sample preparation</u> - Library preparation for RRBS was performed using Premium RRBS Kit (Diagenode) according to manufacturer’s instructions. Briefly, 100ng of DNA was used for each sample, which was enzymatically digested, end-repaired and ligated with an adaptor. Subsequently, 8 samples with different adaptors were pooled together and subjected to bisulfite treatment. After purification steps following bisulfite conversion, the pooled DNA was amplified with 9-14 cycles of PCR and then cleaned up with Ampure XP beads. Quantification of libraries were performed using real-time qPCR (Thermo Fisher). Libraries were sequenced single end 101bp on Hiseq2500 (Illumina).

#### Sequencing alignments, QC, and fingerprinting

<u>DNA alignments</u> - DNA alignments for whole genome (WGS) and exome sequencing was done using bwa-mem (version 0.7.15-r1140) (Fleshner and Chernett, 1997) with -M -t 12 argument and against CanFam3.1 reference genome from UCSC, https://genome.ucsc.edu/cgi-bin/hgGateway?db=canFam3 (md5: 112bc809596d22c896d7e9bcbe68ede6). For each sample, fastq files were aligned per read group and then merged using Picard tools (v2.18.0, http://broadinstitute.github.io/picard) *SortSam* command to make an interim bam file. Final, analysis-ready bam file per sample – tumor and normal bam, if available – was created by series of steps following best practices guidelines from GATK4 (version 4.0.8.1) (DePristo et al., 2011), namely *MarkDuplicates*, *Indel Realignment*, and *Base Quality Score Recalibration* (BQSR). Alignment QC metrics were calculated using GATK4 *DepthOfCoverage* (for WGS) and *CollectHSMetrics* (for exome) as well as Qualimap (version 2.2.1) (Okonechnikov et al., 2016) *bamqc* for merged bam files. Coverage statistics were also based on regions of interest (ROIs) which consisted of exonic region-level annotations for biotypes: protein-coding gene, microrna, lincrna, and pseudogene from Ensembl gene annotations for canine genome (v91 and higher). We flagged samples as failed QC if merged bam file has a genome-wide coverage of < 30% or > 75% of ROIs have 30% or lesser coverage. Accordingly, three samples (of three cases) failed QC step and they were removed from all analyses. Note that 77 cases in patient tissues and samples section represent all cases which passed QC at WGS, exome, RNA-seq, and methylation level data preprocessing. <u>RNA alignments</u> - Raw fastq files from paired-end RNA-seq assay for 41 tumor samples and 3 matched normal tissue samples were first preprocessed through *fastp* (version 0.19.5) (Chen et al., 2018) to perform read-based quality pruning, including adapter trimming. Resulting fastq files were then used as input for *kallisto quant* (version 0.45.0) – a pseudoalignment based method to quantify RNA abundance at transcript-level in transcripts per million (TPM) counts format. We then used *sleuth* R package (version 0.30.0) (Pimentel et al., 2017) to output model-based, gene-level normalized TPM matrix which was also corrected for potential batch effects due to RNA-seq data derived from two sequencing core facilities and tissue archival (snap-frozen vs FFPE). Detailed workflow, including command-line parameters for model fitting are in extended methods section. <u>RRBS alignments</u> - Raw fastq files from RRBS assay for 45 tumor samples were processed through FastQC (version 0.11.7, https://www.bioinformatics.babraham.ac.uk/projects/fastqc) and Trim Galore (version 0.5.0, https://github.com/FelixKrueger/TrimGalore) for quality control, filtering low quality base calls, and adapter trimming. Trimmed reads were then mapped to a bisulfite converted reference genome (canFam3.1, obtained from Ensembl release 85) using the Bismark Bisulfite Mapper (v0.19.1) with the Bowtie2 short read aligner (v2.2.3) (Krueger and Andrews, 2011), allowing for one non-bisulfite mismatch per read. Cytosine methylation calls were made for the mapped reads using the Bismark methylation extractor (version 0.19). The resulting methylation values were obtained as β-values, calculated as the ratio of methylated to total reads at a given CpG site. <u>DNA fingerprinting</u> – DNA fingerprinting for each of WGS and exome tumor-normal and tumor-only bam files was done using *NGSCheckMate* tool (version 1.0.0) (Lee et al., 2017). Germline snps in protein-coding regions was used as a variant reference panel to allow simultaneous fingerprinting of WGS and exome libraries. *NGSCheckMate* does sample pairing QC based on shared germline variants found in samples (tumor and normal tissue from the same patient) and also model difference between samples (or libraries) based on sequencing depth-dependent variation in allelic fraction of reference variants. Fingerprint results for WGS and exome samples from 77 canine glioma did not show mixture of tumor-normal or cross-patient sample contamination (See **Figure S1I**).

#### Somatic variant calling

Somatic variant calls were called on the merged whole genome and exome bam files using three callers: GATK4 (version 4.0.8.1) (McKenna et al., 2010) Mutect2 (Cibulskis et al., 2013), VarScan2 (version 2.4.2), and LoFreq (version 2.1.3.1) (Wilm et al., 2012). Matching and fingerprint validated WGS and exome files per sample were merged using Picard tools (v2.18.0, http://broadinstitute.github.io/picard), *MergeSamFiles* command. Three somatic callers were then run in either paired tumor – matched normal (n=57) or tumor-only (n=20) mode. Mutect2 was first run in panel-of-normals (PON) mode using 57 matched normal samples. Resulting PON file was used for calling somatic variant calls using Mutect2 in both, paired and tumor-only mode along with options: --germline-resources 58indiv.unifiedgenotyper.recalibrated_95.5_filtered.pass_snp.fill_tags.vcf.gz –af-of-alleles-not-in-resource 0.008621. Tumor-only Mutect2 mode was run using default arguments and paired Mutect2 calls had following arguments: --initial-tumor-lod 2.0 -- normal-lod 2.2 --tumor-lod-to-emit 3.0 --pcr-indel-model CONSERVATIVE. Throughout the process of using GATK4 based tools, including Mutect2, we followed best practices guidelines (DePristo et al., 2011) where practical for canine genome, e.g., in contrast to human genome, population level resources are limited for canine genome. VarScan2 paired mode was run with a command: *somatic* and arguments: --min-coverage 8 --min-coverage-normal 6 --min-coverage-tumor 8 --min-reads2 2 --min-avg-qual 15 --min-var-freq 0.08 --min-freq-for-hom 0.75 --tumor-purity 1.0 --strand-filter 1 --somatic-p-value 0.05 --output-vcf 1. VarScan2 tumor-only mode was run using command: *mpileup2cns* and arguments: --min-coverage 8 --min-reads2 2 --min-avg-qual 15 --min-var-freq 0.08 --min-freq-for-hom 0.75 --strand-filter 1 --p-value 0.05 --variants --output-vcf 1. LoFreq paired mode was run using command: *somatic* and arguments: --threads 4 --call-indels --min-cov 7 –verbose and tumor-only mode was run using command: *call* and arguments: --call-indels --sig 0.05 --min-cov 7 --verbose -s. Resulting raw somatic calls - single nucleotide variants (SNV) and small insertions and/or deletions (Indels) - from three callers were then subject to filtering based on caller-specific filters and hard filters. Briefly, Mutect2 calls were subject to extensive filtering based on germline risk, artifacts arising due to sequencing platforms, tissue archival (FFPE), repeat regions, etc. See extended methods and https://software.broadinstitute.org/gatk/documentation/article?id=11136). VarScan2 somatic filters were applied as per developer’s guidelines (Koboldt et al., 2013). Hard filters were based upon filtering out variants present in dbSNP and PONs created via GATK4 Mutect2. Filtered somatic calls from three callers (in VCF version 4.2 format) were then subject to consensus somatic calls using SomaticSeq (version 3.1.0) (Fang et al., 2015) in majority voting mode with priority given to Mutect2 filtered (PASS) calls followed by consensus voting based on calls present in VarScan2 and LoFreq filtered calls. Resulting consensus VCF file for 77 cases were finally converted to Variant Effect Predictor (VEP version 91) (McLaren et al., 2016) annotated vcfs and Mutation Annotation Format (MAF, https://docs.gdc.cancer.gov/Data/File_Formats/MAF_Format) using *vcf2maf* utility (https://github.com/mskcc/vcf2maf). Annotated VCFs and MAFs were used for all of downstream analyses.

#### Significantly mutated genes (SMGs) analysis

SMG analysis in canine gliomas (**Figure 1A, 1C, 2A**) with paired tumor-normal samples (n=57) was performed using dNdS (Martincorena et al., 2017) and MuSiC2 (version 0.2) (Dees et al., 2012). We excluded tumor-only cases for being conservative in SMG analysis and minimize false-positives. Also, MuSiC2 required matched normal tissue required matched normal tissue for SMG analysis. Detailed parameters for SMG analysis are in extended methods. Detailed output from both methods in respectively in table S2 and table S3.

#### Cancer hallmark analysis

Cancer hallmarks were defined according to published ten hallmarks (Hanahan and Weinberg, 2011) and one additional hallmark, i.e. epigenetic (Imielinski et al., 2012). A pool of known glioma and pan-cancer driver genes (Ceccarelli et al. 2016; Gröbner et al., 2018; Bailey et al. 2018) were mapped to hallmarks following a previously published computer-assisted manual curation method (Table S4) (Iorio et al., 2018). Total five cohorts were defined, i.e. IDHwt- (n=371), IDHmut-codel (n=169) and IDHmut-noncodel (n=268) subgroups in human adult gliomas (AG) (Ceccarelli et al. 2016), human pediatric glioma (PG) (Gröbner et al. 2016) and canine glioma (CG). For each of the five cohorts coding mutations were mapped to eleven hallmarks and coverage adjusted relative proportions of patients harboring an alteration in a given hallmark were calculated. For comparisons between cohorts a two-sided Fisher’s exact test was applied (Table S5).

#### Telomere length estimation

Telomere length (TL) estimation (**Figures S1C, S1D**) was done using *telomerecat* (version 3.2) (Farmery et al., 2018) tool given it does not assume fixed number of telomeres (per chromosomes) and hence, can better model (Farmery et al., 2018) telomere length in samples with aneuploidy like canine gliomas. Using *bam2telbam* command, we first extracted telomeric reads from WGS bam files of tumor and matched normal samples. Telomere length per tumor and matched normal samples were then estimated using telbam2length command which counts number of telomeric reads containing the sequence “TTAGGG” or “CCCTAA” for two or more times. Additional arguments, -N 10 and -e were used to run length simulation and cohort wise error correction (snap-frozen vs FFPE) for accurate estimation of telomere length. Telomere length for human pediatric and adult tumors were taken from published studies (Barthel et al., 2017; Parker et al., 2012). Gene expression of core telomere pathway genes (*TERT, ATRX, DAXX*) was calculated from processed RNA-seq data (**Figure S1E**).

#### Quantifying somatic mutation rates

Somatic mutations (SNVs and Indels) rate was estimated within coding genes and adjusted based on relative per-base coverage with minimum coverage of 30X in coding regions (**Figure 1D**). Coding mutation rates for human pediatric (n=961) and adult cancers (n=3,800, includes 811 adult gliomas) were taken from published studies (Ceccarelli et al., 2016; Gröbner et al., 2018; Ma et al., 2018).

#### Somatic copy number segmentation and GISTIC2 based significance

Somatic copy-numbers were called for paired tumor-normal cases (n=56) using HMMCopy tool (version 1.22.0) using author’s recommendations. In brief, GC counts and mappability files for CanFam3.1 genome were generated with 1000 bp window size. Read counts for each of tumor and normal bam files were generated using 1000 bp window size. Resulting count, mappability and count files were feed into HMMCopy algorithm (http://bioconductor.org/packages/release/bioc/html/HMMcopy.html) and segmentations were called using Viterbi algorithm. Segmented copy number calls were used to generate Integrated Genome Viewer (IGV) copy-number plots and GISTIC2 (version 2.0.22) based somatic copy number significance (Mermel et al., 2011), including calling gene-level deep deletions, loss-of-heterozygosity (LOH), and amplifications (**Figure 2A**) as well as inferring aneuploidy metrics (**Figure 2B, 2C, 2D**). Segmented copy number for pediatric gliomas (n=53) were called by using cloud-based TitanCNA workflow (https://dxapp.verhaaklab.com/dnanexus_ngsapp). Segmented copy number for adult gliomas were derived from SNP6 based platform from the TCGA Broad Firehose platform (version stddata 2016_01_28) with following download urls: http://gdac.broadinstitute.org/runs/stddata_2016_01_28/data/GBM/20160128/gdac.broadinstitute.org_GBM.Merge_snp_genome_wide_snp_6_broad_mit_edu_Level_3_segmented_scna_minus_germline_cnv_hg19_seg.Level_3.2016012800.0.0.tar.gz and http://gdac.broadinstitute.org/runs/stddata_2016_01_28/data/LGG/20160128/gdac.broadinstitute.org_LGG.Merge_snp_genome_wide_snp_6_broad_mit_edu_Level_3_segmented_scna_minus_germline_cnv_hg19_seg.Level_3.2016012800.0.0.tar.gz Only primary tumor cases from TCGA GBM (n=577) and TCGA LGG (n=513) cohort were used for downstream analyses, i.e., pathway analysis (**Figure 1C**) and aneuploidy metrics (**Figure 2B, 2C, 2D**).

#### Allele specific copy-number analysis

We derived allele-specific copy numbers and copy-number based clonality inference (including purity and ploidy estimates) using TitanCNA algorithm (version 1.19.1) (Ha et al., 2014). *Snakemake* (version 5.2.1) based workflow(Koster and Rahmann, 2018) was implemented using default arguments and genome-specific germline dbSNP resource (see under extended methods) (https://github.com/gavinha/TitanCNA/tree/master/scripts/snakemake) for WGS paired tumor-normal samples from 56 canine patients. For pediatric gliomas (n=53) and adult gbms with WGS data (n=23), allele-specific copy-number calls were used from TitanCNA workflow. Allele-specific copy-numbers were used for mutational signature and molecular timing analysis (**Figure 3**).

#### Aneuploidy metrics

The simplest metric of aneuploidy was computed by taking the size of all non-neutral segments divided by the size of all segments. The resulting aneuploidy value indicates the proportion of the segmented genome that is non-diploid. In parallel, an arm-level aneuploidy score modeled after a previously described method was computed (Taylor et al., 2018). Briefly, adjacent segments with identical arm-level calls (−1, 0 or 1) were merged into a single segment with a single call. For each merged/reduced segment, the proportion of the chromosome arm it spans was calculated. Segments spanning greater than 80% of the arm length resulted in a call of either −1 (loss), 0 (neutral) or +1 (gain) to the entire arm, or NA if no contiguous segment spanned at least 80% of the arm’s length. For each sample the number of arms with a non-neutral event was finally counted. The resulting aneuploidy score is a positive integer with a minimum value of 0 (no chromosomal arm-level events detected) and a maximum value of 38 (total number of autosomal chromosome arms – given all of canine chromosomes are either acrocentric or telocentric).

#### Syntenic regions and clustering based on aneuploidy metrics

Shared syntenic regions between CanFam3.1 and hg19 reference genome were downloaded from Ensembl BioMart (version 94) database using orthologous mapped Ensembl gene ids. Arm-level synteny was based on arm-level aneuploidy scores of shared syntenic regions in the respective, canine and human genomes. Hierarchical clustering of syntenic arms was then carried out for each of canine, human pediatric and adult cohort (**Figure 2C**).

#### Mutational signature analysis

Mutational signature analysis was performed in two-parts. First, de-novo signatures (**Figure 3B**) were constructed for canine (n=77), human pediatric (n=53) and adult cohort (n=23) using somatic snvs from whole-genome data. Signatures were constructed using non-negative matrix factorization (*nmf* R package, version 0.21.0) with brunet approach and 100 iterations with expected range of signatures between 2 to 10. Optimal signatures were then selected using *nmfEstimateRank* function to match number of de-novo signatures (clusters) – 1 where inflection point for cophenetic correlation coefficient was observed. Accordingly, three de-novo signatures were found in canine and human pediatric gliomas while five in adult glioblastoma cohort. In the second part, known human mutational signatures from COSMIC (v2, n=30) and published pediatric cancer signature from two studies, T1 to T11 (Ma et al., 2018) and P1 (Gröbner et al., 2018) were pooled together and used to deconvolute (*MutationalPattern* R package, version 1.6.2) mutational trinucleotide context (n=96) from somatic snvs in each of three cohorts. Somatic ultra-hypermutation cases from pediatric (n=3) and adult cohort (n=1) were excluded from signature analysis. Cosine similarities of known signatures with de-novo signatures was then calculated and clustered using hierarchical clustering (**Figure 3B**). Absolute and relative contribution of known signatures per sample was then quantified using *fit_to_signatures* function which finds the linear combination of signatures that closely resembles 96 context based mutational matrix by solving the nonnegative least-squares constraints problem. We then selected top contributing signatures per cohort based on signatures which contributed per sample to higher than 3^rd^ quartile of median value of each signature’s contribution (rowMedian) per cohort (**Figure S3A**). Top contributing signatures were further calculated using *outlier profling* on canine patients showing highest mutational load (>3^rd^ quartile of median coding mutation rate per megabase) and plotted in **Figure 3A**. Signatures contributing to driver mutations (**Figure 3C**) were calculated based on first getting relative proportion of trinucleotide context per snv and then finding known signatures with maximum value for the same trinucleotide context. Known signatures were combined to a single group where they are shown in literature as potential underlying process, e.g., aging group is associated with COSMIC signature 1 and 5, and show significant cosine similarity (> 0.9) with pediatric signatures T1 and T4, respectively. Table S6 provides mapping between signature and known/proposed mechanisms, if any.

#### Molecular timing analysis and natural history of tumors

Probabilistic estimation of relative timing of driver mutations (among 73 observed somatic snvs in cancer driver genes) was based on existing methods (Gerstung et al., 2017; Jolly and Van Loo, 2018) with several steps carried out using Palimpsest R package (version 1.0.0; https://github.com/FunGeST/Palimpsest) (Shinde et al., 2018) and custom R scripts based on published approach(McGranahan et al., 2015): First step involved categorizing somatic drivers into clonal vs subclonal events using estimated cancer cell fraction (CCF) which is estimated fraction of cancer cells with a somatic snv. CCF per somatic snv was a product of variant allelic fraction (VAF) of a somatic snv, adjusted by local copy number of gene locus and whole tumor sample (ploidy) as well as purity estimate (tumor cell content) inferred from TitanCNA algorithm (Detailed under copy number estimation section above). A clonal (early) vs subclonal (late) mutation was then classified based on upper boundary of CCF was above 0.95 (clonal) or not (subclonal). Second, we timed copy number gain and copy-neutral LOH regions based on VAF of somatic snvs in these copy regions, i.e., early mutations prior to copy gain will have higher VAF relative to VAF of late mutations after copy gain. Third, we ordered mutations in four sequential categories: early clonal, early subclonal, late clonal, and late subclonal. We note here that early subclonal and late clonal categories are result of underlying parallel and/or convergent evolution of multiple clones (Venkatesan and Swanton, 2016) and/or a technical limitation (given ∼60X depth of merged bam files and lack of spatial sequencing data) in resolving polyclonal structure of a tumor sample (Deshwar et al., 2015). We then tally frequency of each of these four categories per somatic driver mutation and get the average frequency of each category per driver mutation at cohort (canine, pediatric, adult) level. These average frequencies are converted to winning tables, similar to sports statistics where each driver mutation competes with remaining driver mutations with winning being an early somatic event based on order of events using clonality (Jolly and Van Loo, 2018) (step 3). Finally, a winning table is then passed to Bradley-Terry model (*BradleyTerryScalable* R package, version 0.1.0.9000; https://cran.r-project.org/web/packages/BradleyTerryScalable/vignettes/BradleyTerryScalable.html) to estimate winning probability (driver event being an early event) based on a Bayesian maximum a posteriori probability (MAP) estimate. Resulting winning probability per driver mutation is subtracted from 1 to plot multiple density plots (*ggridges* R package, version: 0.5.1.9000) with X-axis now showing a probability of event being a late event (**Figure 3D**). We note that density plots are based on kernel density estimates and thus, may extend their tails (probability distribution) beyond 1 or less than zero (https://serialmentor.com/dataviz/histograms-density-plots.html).

#### Class prediction using methylation data

To compare the methylation patterns of human and canine glioma, the LIBLINEAR library was used to fit an L2-regularized logistic regression classifier. Model training and validation was performed on the human glioma samples and normal controls in the GSE109381 dataset (Capper et al., 2018), with the methylation status of CpGs located in regions of the human genome orthologous to canine CpG islands used to predict DNA methylation-based subtypes of glioma. The methylation categories designated as regression outcome variables were derived from the World Health Organization classification of gliomas: IDH-wildtype adult glioma, IDH-mutant, 1p/19q-intact adult glioma, IDH-mutant, 1p/19q-codeleted adult glioma, adult normal control, pediatric glioma, and pediatric normal control. After model fitting, the logistic regression classifier was applied to the canine samples, using the β-values of CpGs orthologous to the selected 11,495 Illumina 450K probes as input data. For classifier CpG sites in the canine samples with no methylation observations, β-values were predicted using the DNA module of the DeepCpG algorithm, a deep learning algorithm that predicts methylation state based on local DNA sequence context (Angermueller et al., 2017). The logistic regression classifier outputs the probability that a sample matches a given methylation category. Category probabilities were calculated for the canine samples, and these probabilities were compared with sample age, anatomical location, tumor grade, tumor purity, and mutation rate (**Figure 4**).

#### CIBERSORT based expression analysis to study tumor microenvironment

Processed RNA-seq expression matrices from canine (n=40; 25 HGG, 14 LGG, 1 unknown grade), adult (n=703; 529 LGG, 174 GBM), and pediatric glioma (n=92; 42 LGG, 50 HGG) were each run as separate jobs into the CIBERSORT webserver (https://cibersort.stanford.edu) and processed in relative mode using the following parameters: Signature Genes: LM22 CIBERSORT default, Permutations run: 100, Quantile normalization disabled (Newman et al., 2015). The resulting cellular fraction tables were then collapsed from 22 cell types into 11 based on lineage, using groupings from a prior publication (Gentles et al., 2015).

## Supplementary Figures

**Figure S1A:**
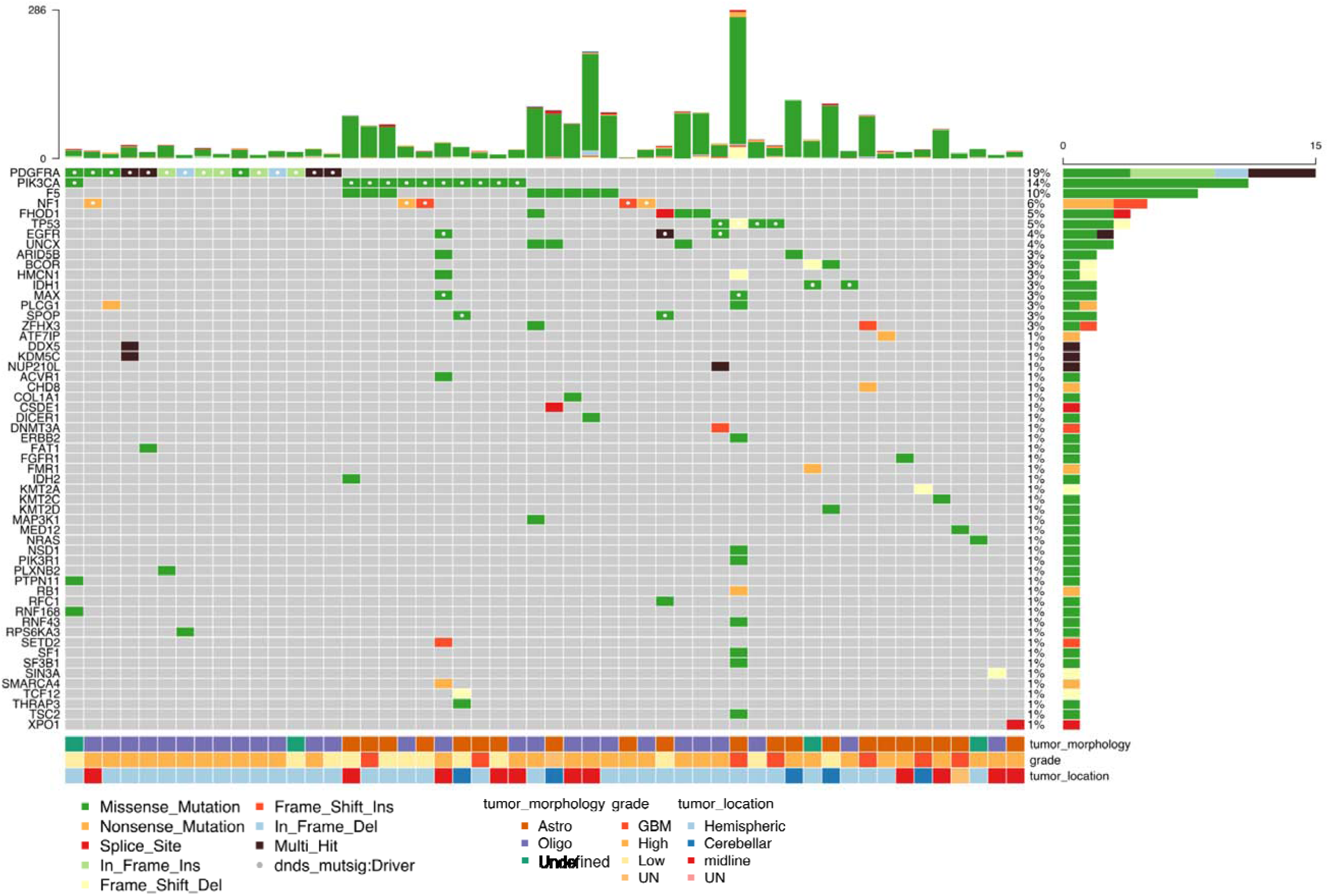
Somatic coding mutations in COSMIC cancer genes (n = 78) *Oncoprint heatmap showing c*olumns show each of 52 of 77 mutated dog patients. Each row is a known cancer gene in COSMIC database. Colored boxes represent type of somatic variant. Top bar plot shows patient-wise frequency of somatic variants and right sided barplot shows driver-wise frequency of somatic variant types. Bottom annotations show relevant patient-specific annotations.

**Figure S1B:**
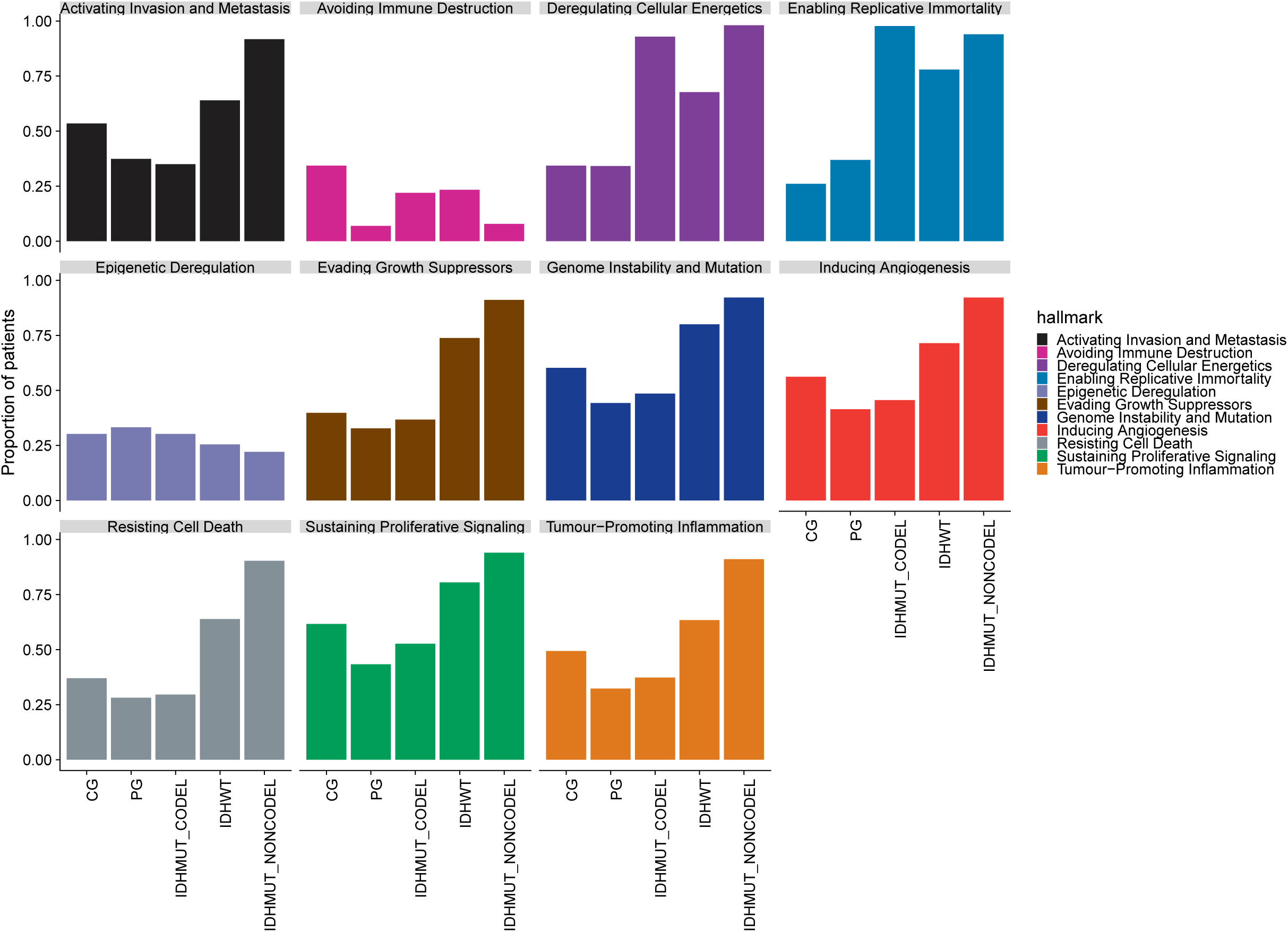
Hallmark enrichment of coding mutations in cancer driver genes across human adult (*IDH*wt, *IDH*mut-codel, *IDH*mut-noncodel), pediatric (PG) and canine (CG) glioma. Eleven hallmarks are presented. Y-Axis represents proportion of patients in the respective cohort harboring mutations in mutations in the hallmarks (different colors). Two-sided Fisher’s exact test used for comparison of proportions between cohorts. Respective p-values are presented in Table S5.

**Figure S1C-E:**
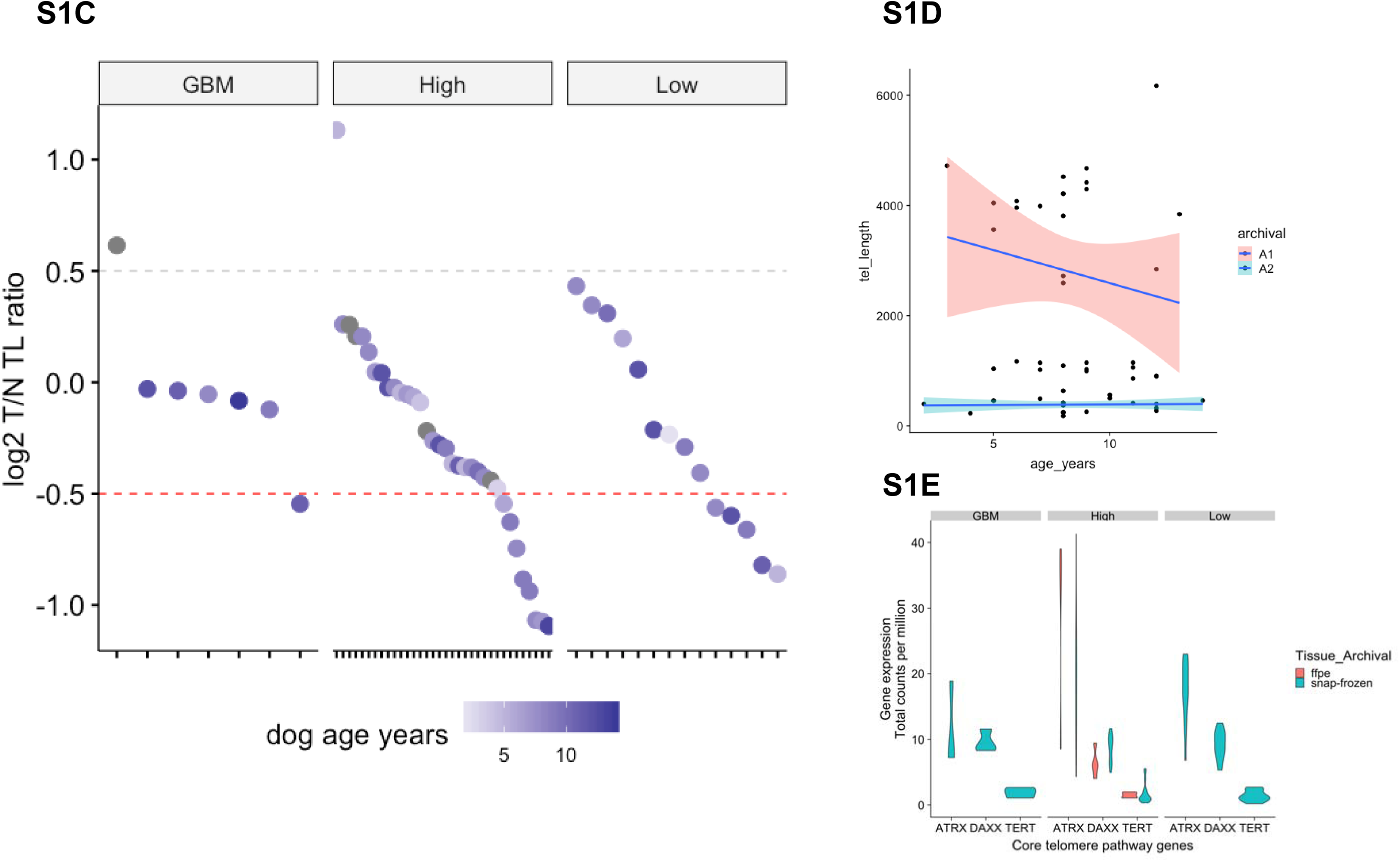
Telomere length estimation in canine gliomas. **C)** Telomere length (TL) estimation (see Methods) for paired tumor-normal cases from patients with canine gliomas (n=55). Remaining two cases had errored estimates and were excluded. Y-axis shows TL as log2 ratio of estimated length of telomere in tumor / TL in normal. Cases are split by tumor grade. TL ratio from pediatric and adult gliomas were compared from Parker et al. 2012 and Barthel et al. 2017, respectively (see Methods). **D)** Scatter plot with spline smoothing showing correlation of telomere length from the matched normal samples (n=55) with dog age (years) and stratified by tissue archival (FFPE or snap-frozen). No significant correlation was observed in both, snap-frozen and FFPE cohorts. **E)** Gene expression as total count per million RNA copies of core telomere pathway genes (*TERT*, *DAXX*, *ATRX*) in canine gliomas (n=40). Violin plots shows expression of these genes factored by tissue archival (FFPE or snap-frozen) and tumor grade (GBM, High vs low grade). No significant difference in expression was observed in among tumor grades.

**Figure S1F-G:**
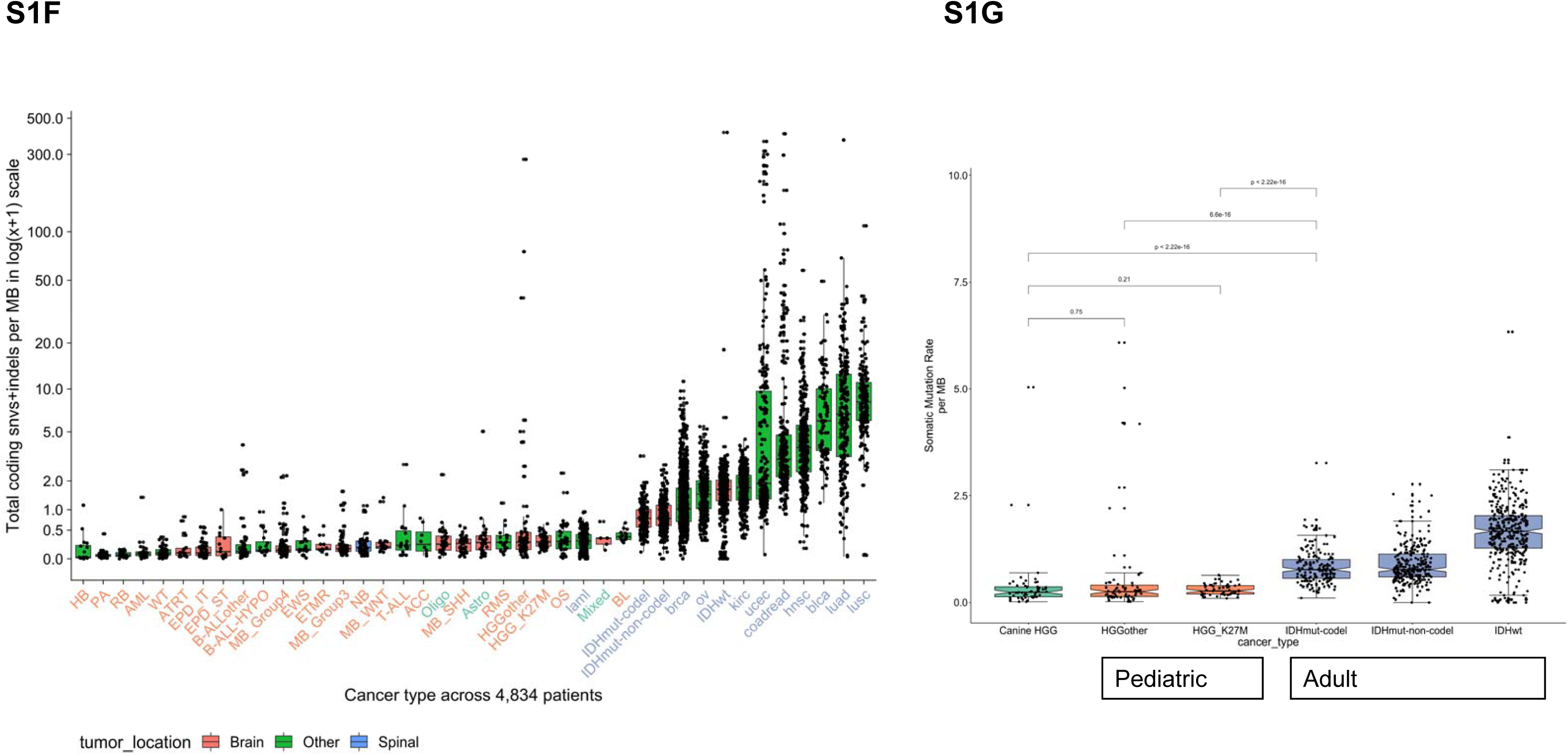
Somatic mutation rate across canine and human pediatric and adult tumors. **F)** Boxplot showing somatic mutation rates as coding mutations per megabase in log1p scale. X-axis shows each of different types of pediatric brain tumors (Grobner 2018), canine glioma (Oligo, Astro, Undefined), and adult GBM (right end of the plot). Tumors are sorted in ascending order by increasing mutation rate. **G)** Boxplot with pairwise differences in somatic mutation rates, including Wilcoxon p-values for canine (n=53) and human pediatric high-grade (57 K27M midline and 64 of other etiology) gliomas vs adult gliomas stratified by IDH1/2 mutations and 1p/19q codeletion (371 IDHwt, 268 IDHmut-non-codel, and 169 IDHmut-codel).

**Figure S1H:**
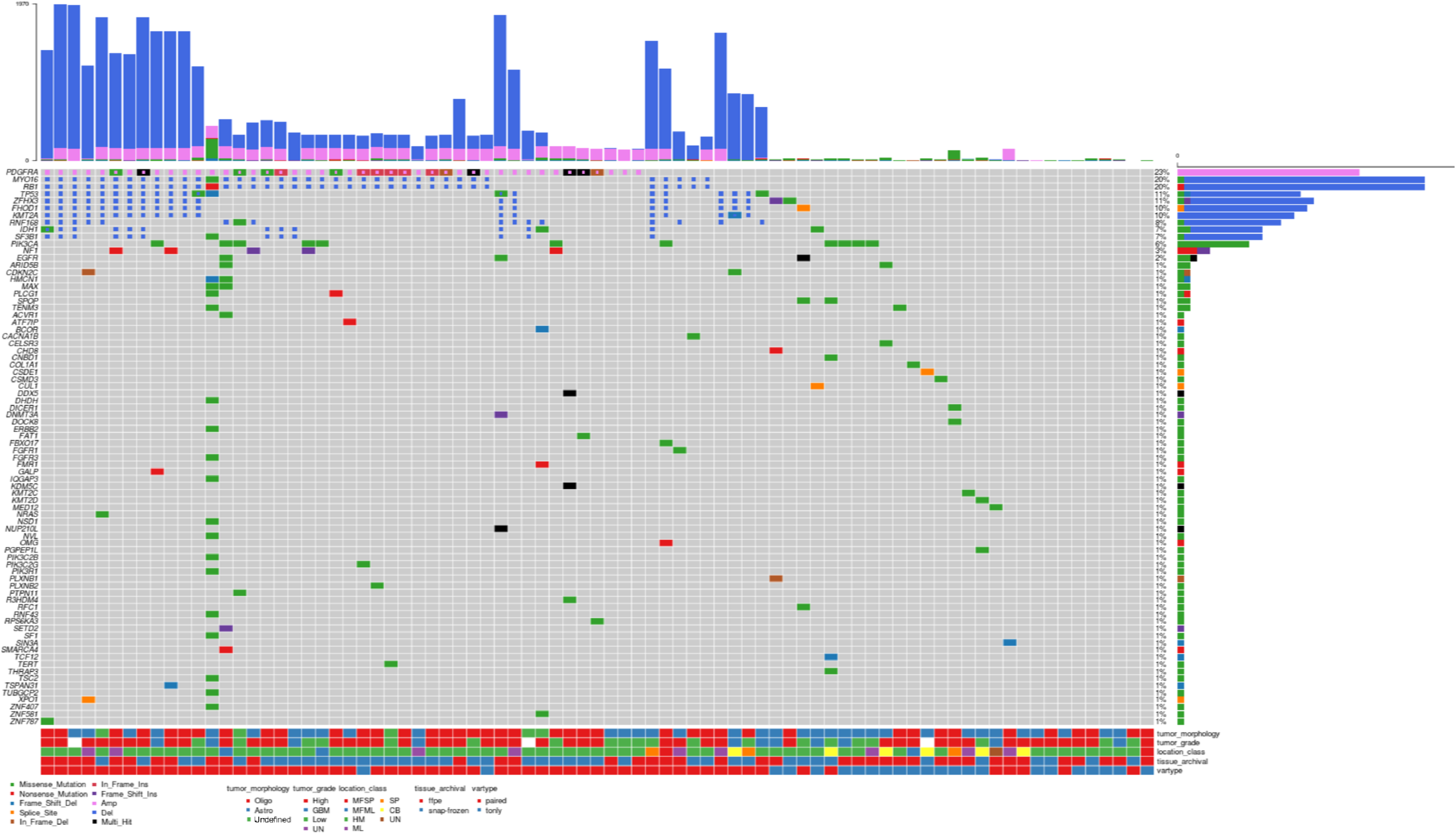
Somatic coding mutations, including copy number aberrations (SCNA) in COSMIC cancer genes (n = 78) *Oncoprint heatmap showing c*olumns show all of 81 dog patients (including 4 blacklisted cases) to show complete somatic landscape among cancer genes in COSMIC database. Each row is a known cancer gene in COSMIC database. Colored boxes represent type of somatic variant, including SCNAs (Amplifications and Deletions). Top bar plot shows patient-wise frequency of somatic variants and right sided barplot shows driver-wise frequency of somatic variant types. Bottom annotations show relevant patient-specific annotations.

**Figure S1I:**
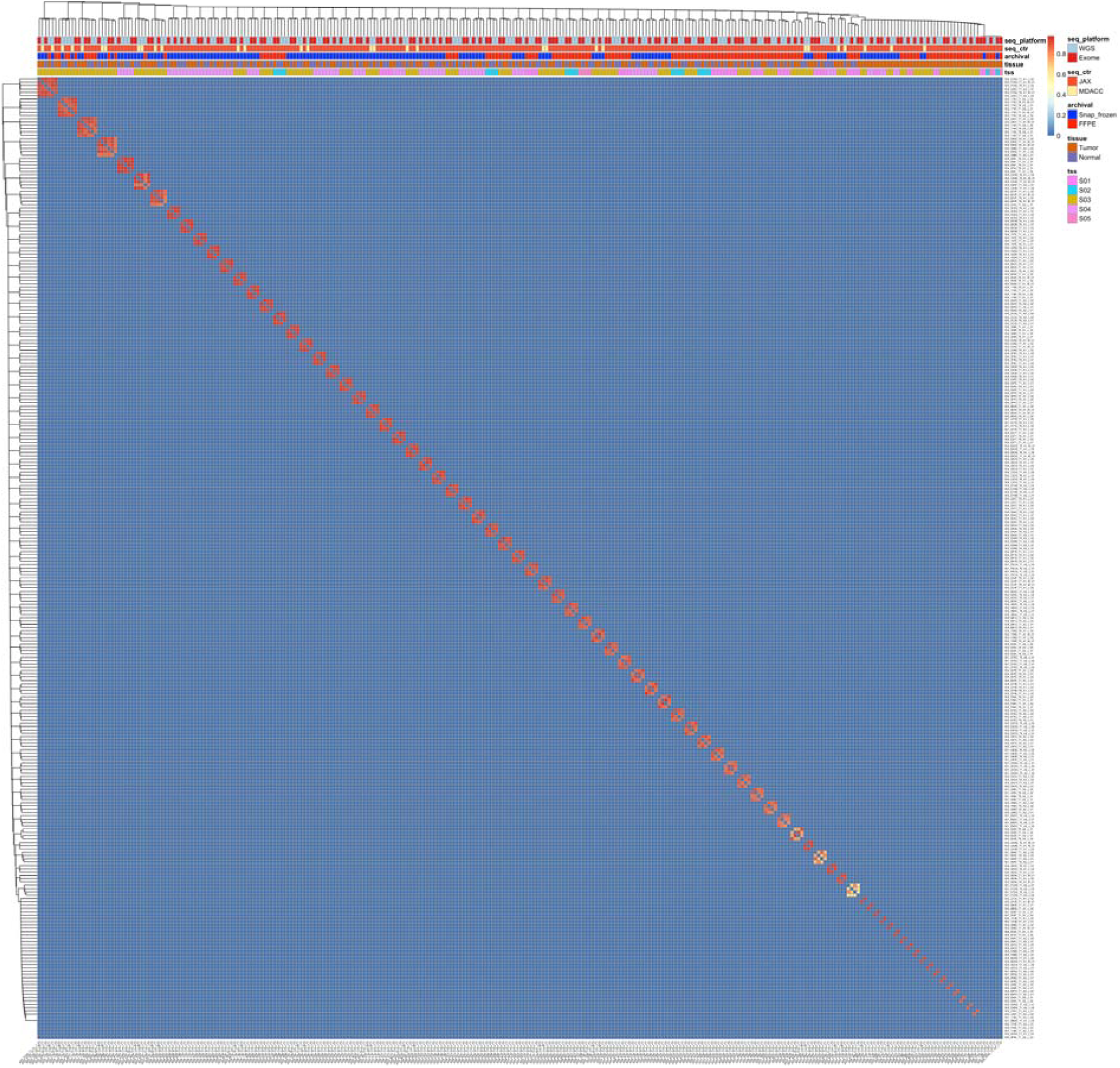
DNA fingerprinting of WGS and exome libraries. This figure related to **STAR methods** section on sequencing alignments, QC, and fingerprinting. Heatmap showing hierarchical clustering (average linkage method) of pairwise correlation matrix between any two sequencing libraries of 77 tumor and 57 normal samples. Higher correlation (red color) between samples suggest sequencing libraries (WGS and exome) originating from the same tissue (tumor or normal) and the same canine patient, and thus grouped into a highly correlating cluster of four samples for paired cases (n=57) or two for tumor-only cases (n=20). Relevant patient-specific annotations are shown in column annotations (top). Correlation matrix is based on the shared germline variants and sequencing depth-based modeling (see Methods for details). TSS: Tissue Source Site

**Figure S2A:**
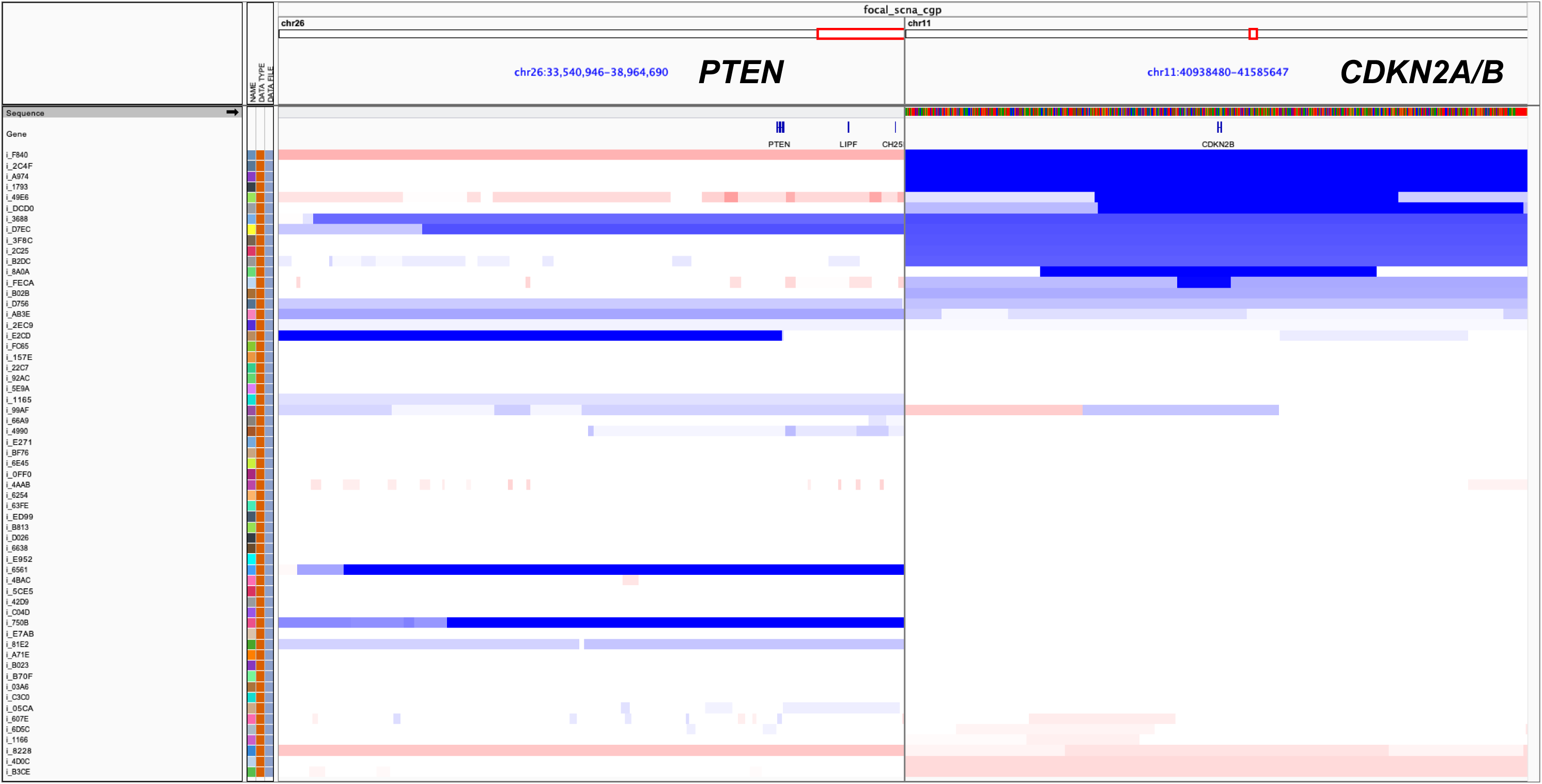
Focal deep deletions in tumor suppressor genes. Integrated Genomics Viewer (IGV) plot showing focal deep or homozygous (GISTIC2 based peak of < −1.1) deletions in *PTEN* (n=3) and *CDKN2A/B* (n=8) regions. Total 11 / 56 (of 57 paired tumor-normal cases; one sample was blacklisted for copy number analysis) dog patients have deep or homozygous deletions in these two genes.

**Figure S2B-C:**
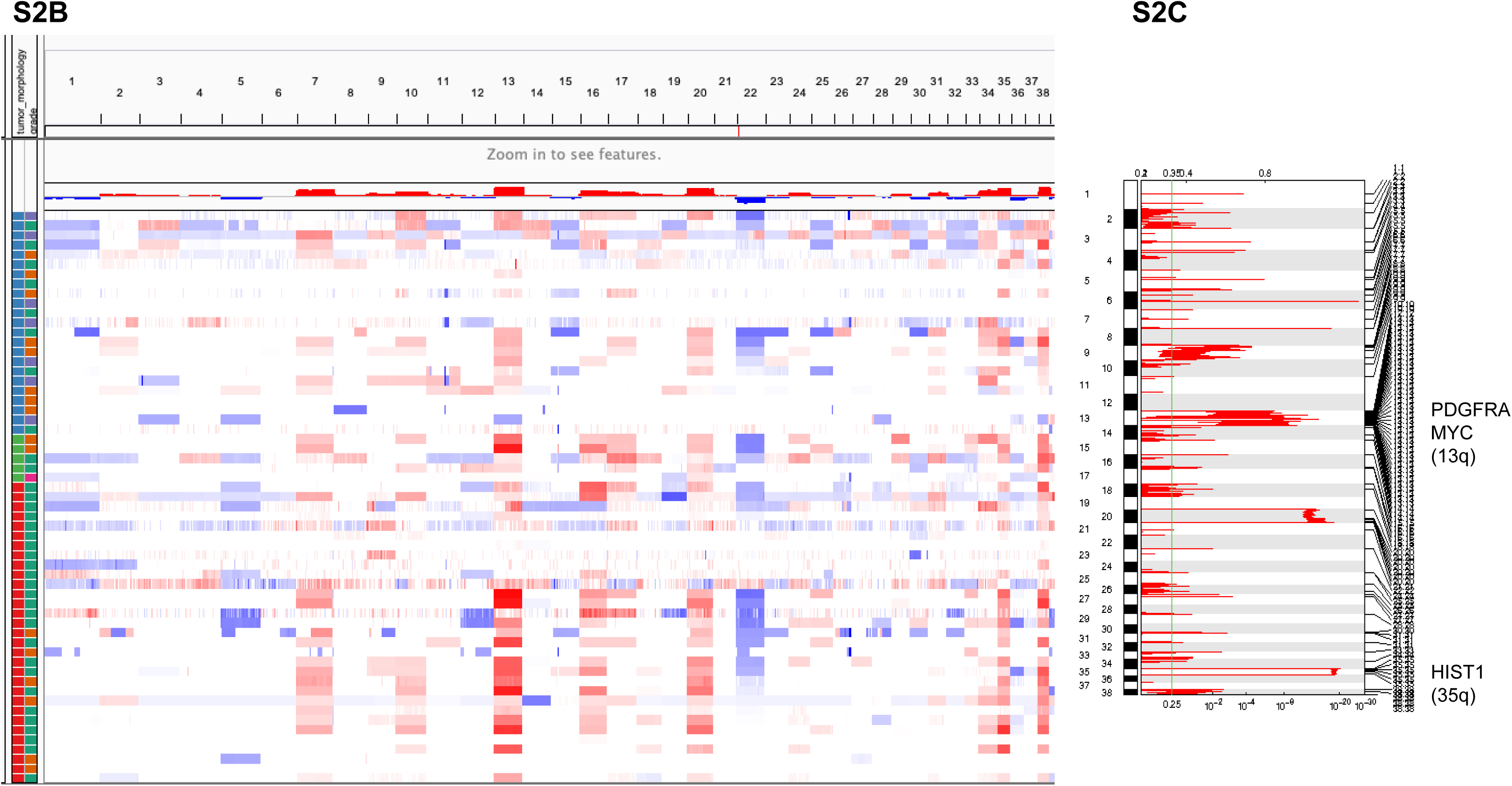
Genome wide somatic copy number alterations. **B)** WGS inferred DNA copy number on paired cases of canine gliomas (n=56 of 57 paired tumor-normal cases; one sample was blacklisted for copy number analysis). IGV plot showing genome-wide copy number changes (red is amplification, blue is deletion) for each of 38 autosomes in 56 dog patients (in rows). Corresponding segmented data is given in table S4. Plot shows frequent gain of chromosomes 7, 13, 16, 20, 34, 35, 3 and frequent loss of chromosomes 1, 5, 12, 22, 26. **C)** Plot shows GISTIC2 based significant focal and broad peaks. X-axis shows GISTIC2 inferred p-values with significant broad or arm-level peaks in multiple chromosomal regions and genes, including in shared syntenic regions of known drivers of human pediatric gliomas.

**Figure S3A:**
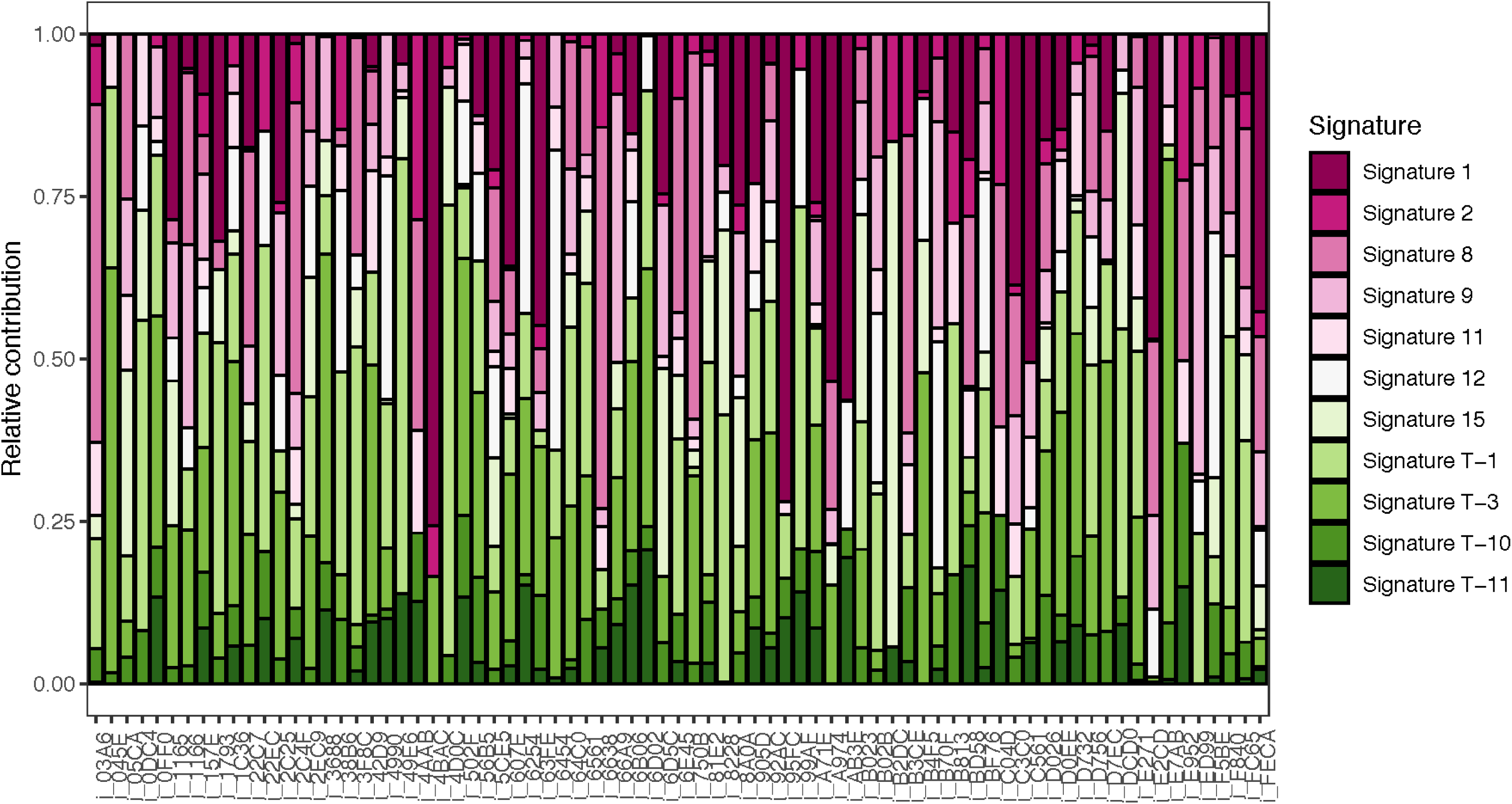
Deconvolution of known human mutational signatures on canine non-hypermutant glioma cases (n=68) Stacked barplots shows relative contribution of known human mutational signatures in individual canine patients. Only top signatures with relative contribution more than third quartile per sample is shown in the plot. Signatures where underlying mechanism is known are colored identically. Table S6 provides mapping between signature and known/proposed mechanisms, if any. Classification of hypermutation vs non-hypermutation was based on outlier profile analysis (see STAR Methods).

**Figure S3B:**
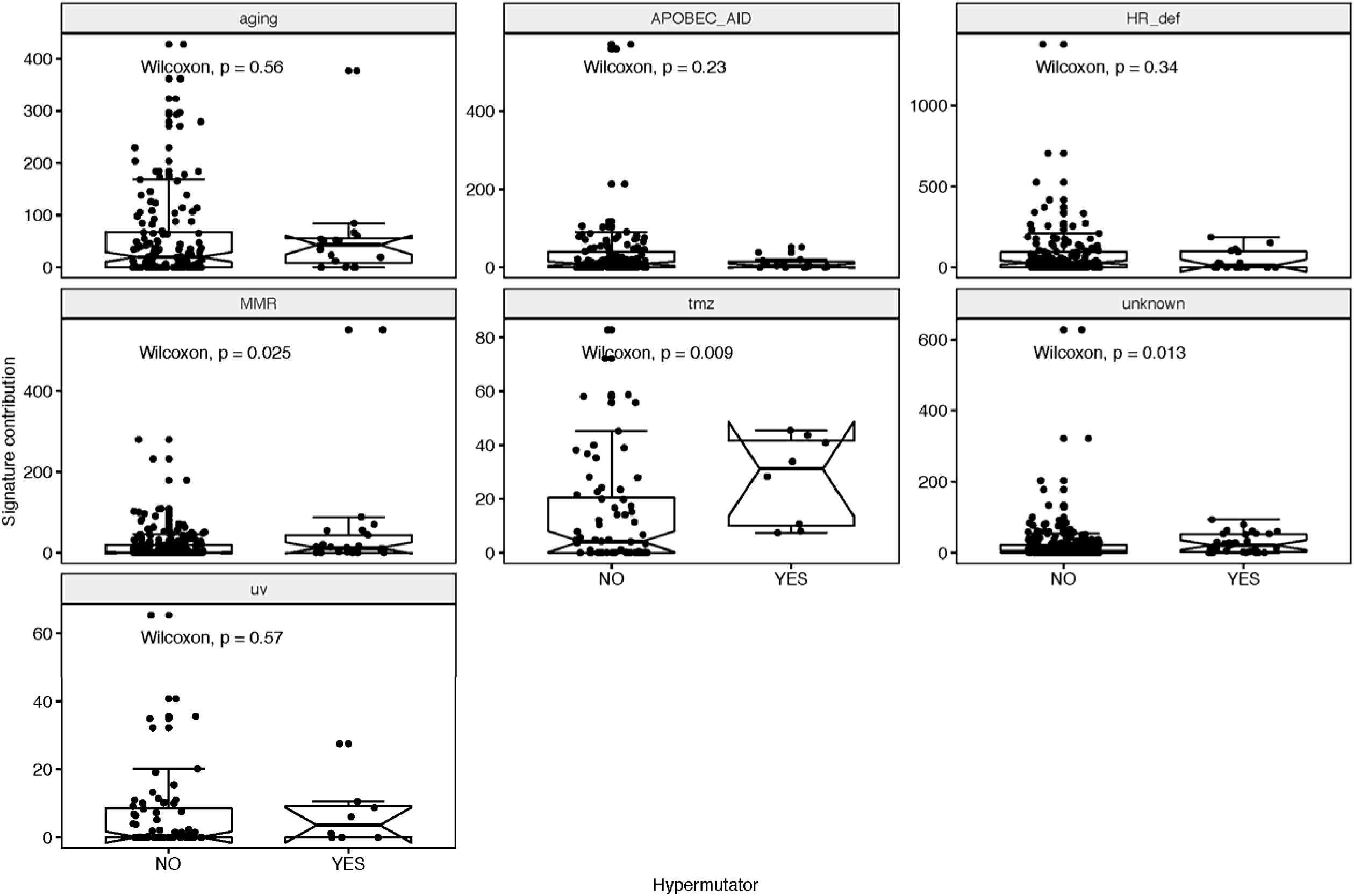
Mutational signature contribution based on hypermutation class. each of seven boxplots with calculated Wilcoxon p-value shows contribution of known human mutational signatures in individual canine patients stratified by mutational load. Classification of hypermutation vs non-hypermutation was based on outlier profile analysis (see Methods). Known human mutational signatures were grouped into seven categories based on known or proposed underlying mechanisms (see Table S6).

**Figure S3C:**
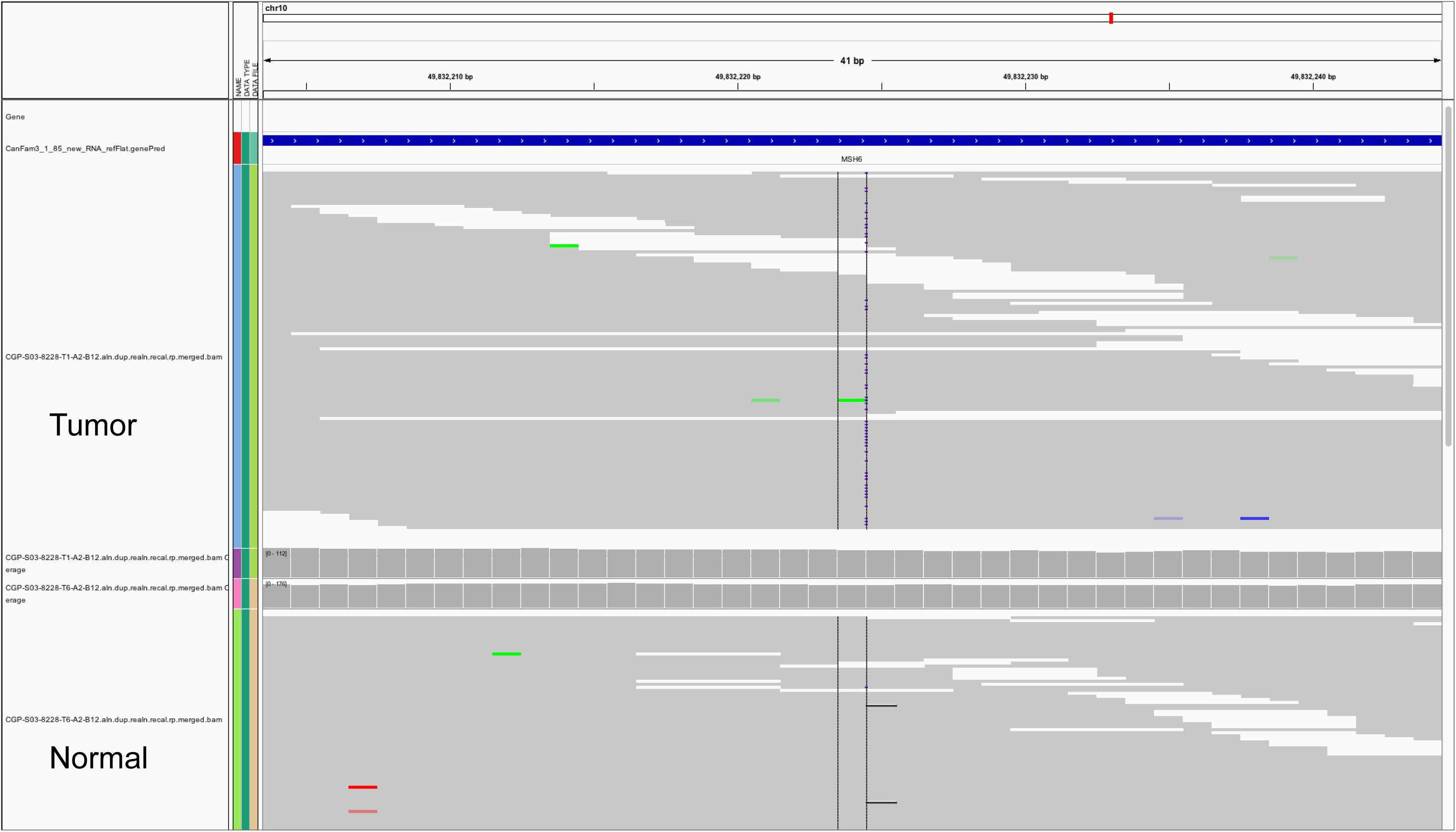
*MSH6* frameshift variant in the hypermutatnt canine patient (i_8228) IGV plot shows sequencing read alignment at *MSH6* locus on canine chromosome 10, showing somatic frameshift variant in reads from tumor sample in contrast to reads from matched normal sample.

**Figure S3D:**
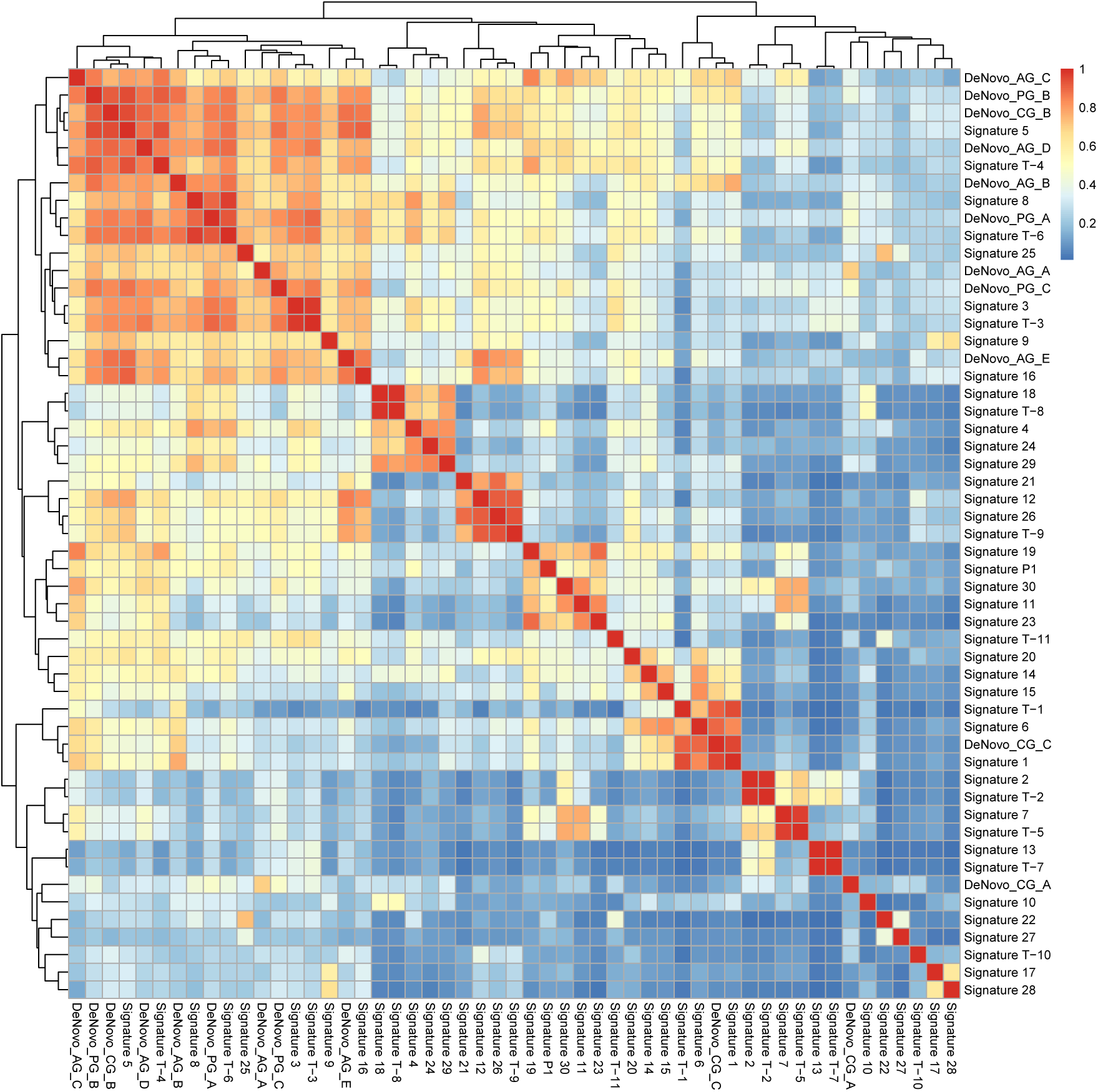
Cosine similarity clustering matrix between known human signatures vs de-novo signatures in canine, adult and pediatric gliomas. See text under molecular life history section and methods for details.

**Figure S3E:**
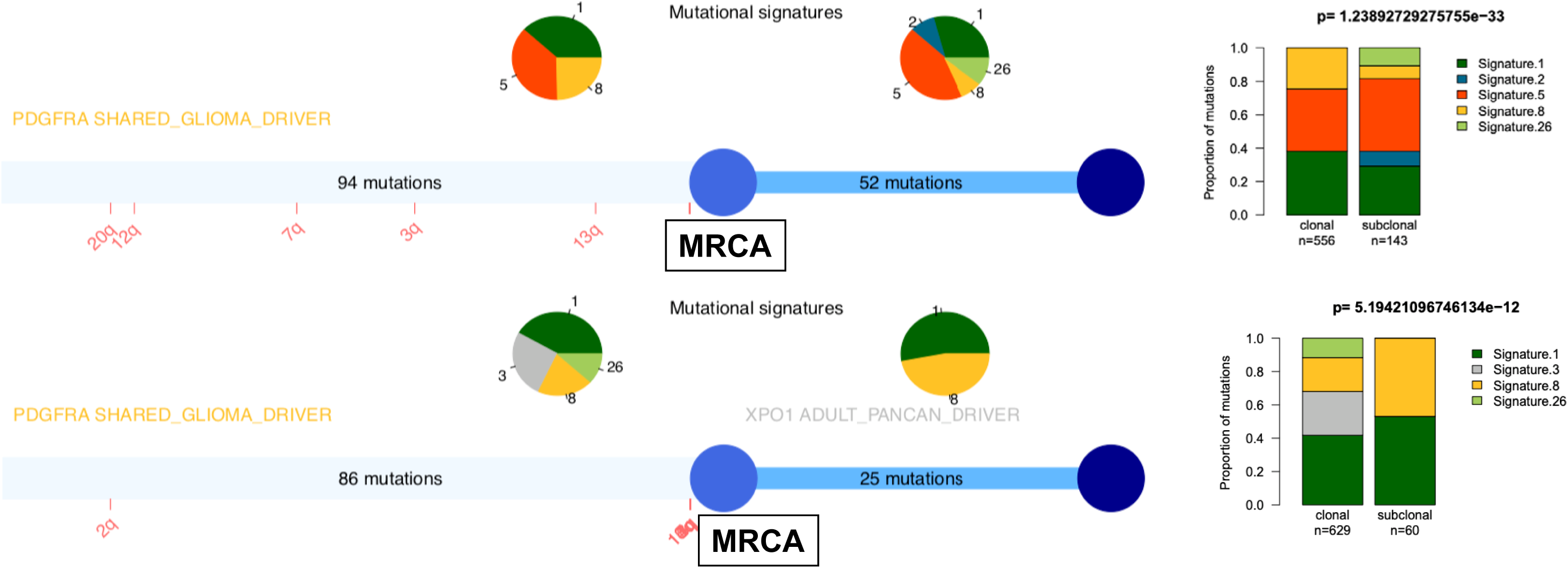
Molecular life history of canine gliomas. Two representative patients of canine gliomas showing driver mutation followed by amplification of *PDGFRA* on chromosome 13q which marks the emergence of most common recent ancestor (MRCA). Case 1 (top) i_5CE5, Case 2 (bottom) i_D026. Relative timing of events is shown for number of chromosomal alterations (only amplifications in these two cases). Gene names are suffixed with their known relevance as a driver in either human adult or pediatric or shared driver (as in these two cases). Gene names are also colored based on type of mutational signature contributing most to mutations in that gene (Signature 8 in these two cases). Pie charts shows relative contribution of mutational signatures before (early) and after (late) emergence of MRCA with respective number of mutations tallied on the straight line. Tallied mutations are coding mutations which can be timed while total number of mutations in these two cases are show in in stacked barplots and colored based on relative contribution of mutational signatures in early vs late phase of inferred tumor evolution.

**Figure S4:**
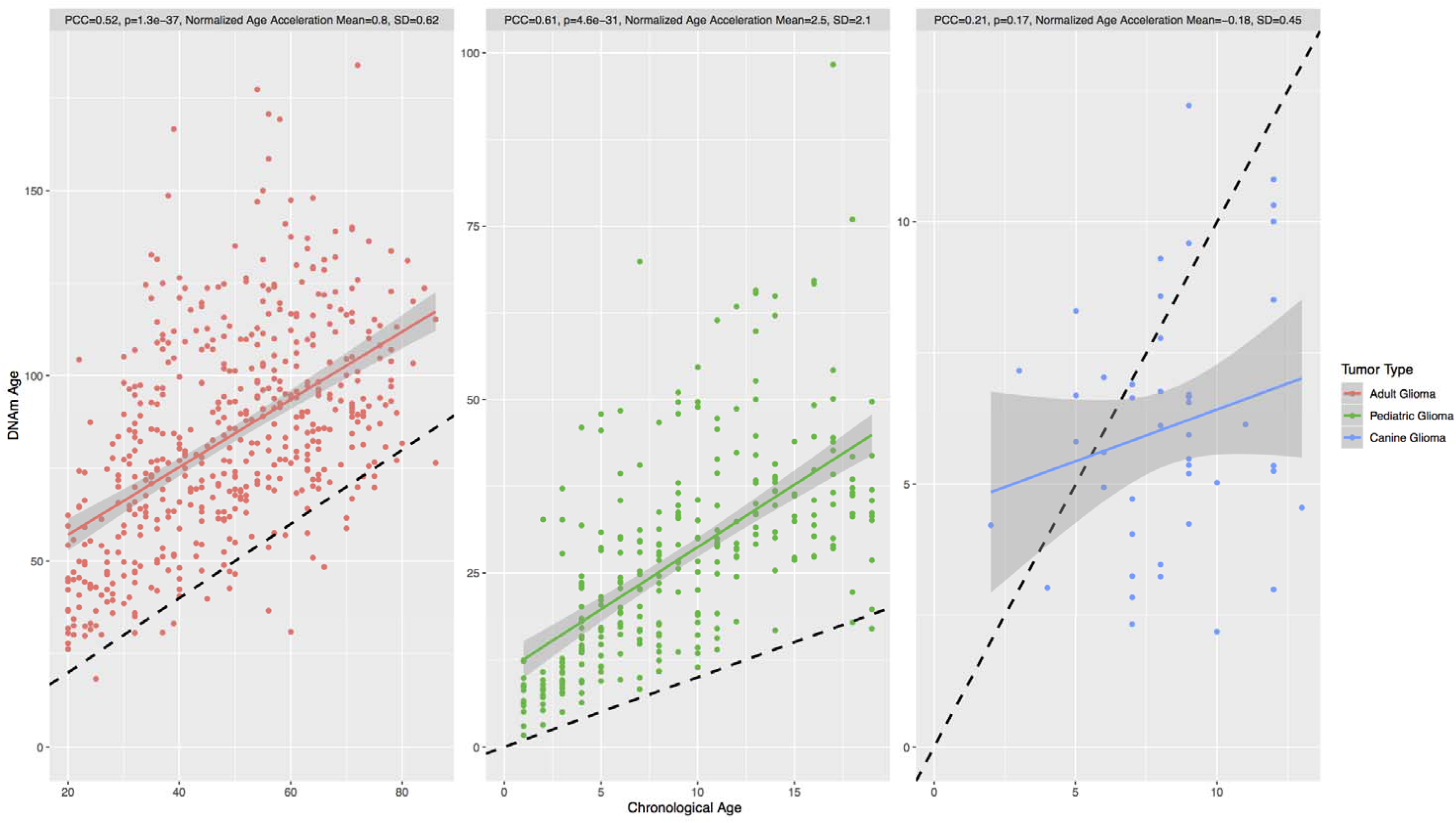
DNA methylation age of human and canine glioma. Scatterplot displaying DNA methylation (DNAm) vs. chronological age of human adult, human pediatric, and canine glioma samples, respectively. The colored line in each plot represents the linear fit from regressing DNAm age on chronological age, and the dotted line represents the linear fit for DNAm age equal to chronological age. The statistics displayed at the top of each subplot are the Pearson correlation coefficient (PCC) of the correlation between DNAm and chronological age, the p-value for the PCC calculation, normalized mean age acceleration, and the standard deviation of normalized age acceleration, respectively.

## REFERENCES

Addissie, S., and Klingemann, H. (2018). Cellular Immunotherapy of Canine Cancer. Vet Sci 5.

Aktipis, C. A., Boddy, A. M., Gatenby, R. A., Brown, J. S., and Maley, C. C. (2013). Life history trade-offs in cancer evolution. Nat Rev Cancer 13, 883–892.

Alexandrov, L. B., Nik-Zainal, S., Wedge, D. C., Aparicio, S. A., Behjati, S., Biankin, A. V., Bignell, G. R., Bolli, N., Borg, A., Borresen-Dale, A. L., et al. (2013). Signatures of mutational processes in human cancer. Nature 500, 415–421.

Alizadeh, A. A., Aranda, V., Bardelli, A., Blanpain, C., Bock, C., Borowski, C., Caldas, C., Califano, A., Doherty, M., Elsner, M., et al. (2015). Toward understanding and exploiting tumor heterogeneity. Nat Med 21, 846–853.

Armitage, P., and Doll, R. (1954). The age distribution of cancer and a multi-stage theory of carcinogenesis. Br J Cancer 8, 1–12.

Bailey, M. H., Tokheim, C., Porta-Pardo, E., Sengupta, S., Bertrand, D., Weerasinghe, A., Colaprico, A., Wendl, M. C., Kim, J., Reardon, B., et al. (2018). Comprehensive Characterization of Cancer Driver Genes and Mutations. Cell 173, 371–385.e318.

Bakhoum, S. F., and Cantley, L. C. (2018). The Multifaceted Role of Chromosomal Instability in Cancer and Its Microenvironment. Cell 174, 1347–1360.

Bakhoum, S. F., Ngo, B., Laughney, A. M., Cavallo, J. A., Murphy, C. J., Ly, P., Shah, P., Sriram, R. K., Watkins, T. B. K., Taunk, N. K., et al. (2018). Chromosomal instability drives metastasis through a cytosolic DNA response. Nature 553, 467–472.

Barthel, F. P., Wesseling, P., and Verhaak, R. G. W. (2018). Reconstructing the molecular life history of gliomas. Acta Neuropathol 135, 649–670.

Blank, H. M., Sheltzer, J. M., Meehl, C. M., and Amon, A. (2015). Mitotic entry in the presence of DNA damage is a widespread property of aneuploidy in yeast. Mol Biol Cell 26, 1440–1451.

Brennan, C. W., Verhaak, R. G., McKenna, A., Campos, B., Noushmehr, H., Salama, S. R., Zheng, S., Chakravarty, D., Sanborn, J. Z., Berman, S. H., et al. (2013). The somatic genomic landscape of glioblastoma. Cell 155, 462–477.

Buque, A., and Galluzzi, L. (2018). Modeling Tumor Immunology and Immunotherapy in Mice. Trends Cancer 4, 599–601.

Capper, D., Jones, D. T. W., Sill, M., Hovestadt, V., Schrimpf, D., Sturm, D., Koelsche, C., Sahm, F., Chavez, L., Reuss, D. E., et al. (2018). DNA methylation-based classification of central nervous system tumours. Nature 555, 469–474.

Ceccarelli, M., Barthel, F. P., Malta, T. M., Sabedot, T. S., Salama, S. R., Murray, B. A., Morozova, O., Newton, Y., Radenbaugh, A., Pagnotta, S. M., et al. (2016). Molecular Profiling Reveals Biologically Discrete Subsets and Pathways of Progression in Diffuse Glioma. Cell 164, 550–563.

Davoli, T., Uno, H., Wooten, E. C., and Elledge, S. J. (2017). Tumor aneuploidy correlates with markers of immune evasion and with reduced response to immunotherapy. Science 355.

Decker, B., Davis, B. W., Rimbault, M., Long, A. H., Karlins, E., Jagannathan, V., Reiman, R., Parker, H. G., Drögemüller, C., Corneveaux, J. J., et al. (2015a). Comparison against 186 canid whole-genome sequences reveals survival strategies of an ancient clonally transmissible canine tumor. Genome Res 25, 1646–1655.

Decker, B., Parker, H. G., Dhawan, D., Kwon, E. M., Karlins, E., Davis, B. W., Ramos-Vara, J. A., Bonney, P. L., McNiel, E. A., Knapp, D. W., and Ostrander, E. A. (2015b). Homologous Mutation to Human BRAF V600E Is Common in Naturally Occurring Canine Bladder Cancer--Evidence for a Relevant Model System and Urine-Based Diagnostic Test. Mol Cancer Res 13, 993–1002.

Dees, N. D., Zhang, Q., Kandoth, C., Wendl, M. C., Schierding, W., Koboldt, D. C., Mooney, T. B., Callaway, M. B., Dooling, D., Mardis, E. R., et al. (2012). MuSiC: identifying mutational significance in cancer genomes. Genome Res 22, 1589–1598.

DeGregori, J. (2017). Connecting Cancer to Its Causes Requires Incorporation of Effects on Tissue Microenvironments. Cancer Res 77, 6065–6068.

Dickinson, P. J., York, D., Higgins, R. J., LeCouteur, R. A., Joshi, N., and Bannasch, D. (2016). Chromosomal Aberrations in Canine Gliomas Define Candidate Genes and Common Pathways in Dogs and Humans. J Neuropathol Exp Neurol 75, 700–710.

Fortunato, A., Boddy, A., Mallo, D., Aktipis, A., Maley, C. C., and Pepper, J. W. (2017). Natural Selection in Cancer Biology: From Molecular Snowflakes to Trait Hallmarks. Cold Spring Harb Perspect Med 7.

Frampton, D., Schwenzer, H., Marino, G., Butcher, L. M., Pollara, G., Kriston-Vizi, J., Venturini, C., Austin, R., de Castro, K. F., Ketteler, R., et al. (2018). Molecular Signatures of Regression of the Canine Transmissible Venereal Tumor. Cancer Cell 33, 620–633.e626.

Gröbner, S. N., Worst, B. C., Weischenfeldt, J., Buchhalter, I., Kleinheinz, K., Rudneva, V. A., Johann, P. D., Balasubramanian, G. P., Segura-Wang, M., Brabetz, S., et al. (2018). The landscape of genomic alterations across childhood cancers. Nature.

Hanahan, D., and Weinberg, R. A. (2011). Hallmarks of cancer: the next generation. Cell 144, 646–674.

Hendricks, W. P. D., Zismann, V., Sivaprakasam, K., Legendre, C., Poorman, K., Tembe, W., Perdigones, N., Kiefer, J., Liang, W., DeLuca, V., et al. (2018). Somatic inactivating PTPRJ mutations and dysregulated pathways identified in canine malignant melanoma by integrated comparative genomic analysis. PLoS Genet 14, e1007589.

Huether, R., Dong, L., Chen, X., Wu, G., Parker, M., Wei, L., Ma, J., Edmonson, M. N., Hedlund, E. K., Rusch, M. C., et al. (2014). The landscape of somatic mutations in epigenetic regulators across 1,000 paediatric cancer genomes. Nat Commun 5, 3630.

Jolly, C., and Van Loo, P. (2018). Timing somatic events in the evolution of cancer. Genome Biol 19, 95.

Khanna, C., Lindblad-Toh, K., Vail, D., London, C., Bergman, P., Barber, L., Breen, M., Kitchell, B., McNeil, E., Modiano, J. F., et al. (2006). The dog as a cancer model. Nat Biotechnol 24, 1065–1066.

Koehler, J. W., Miller, A. D., Miller, C. R., Porter, B., Aldape, K., Beck, J., Brat, D., Cornax, I., Corps, K., Frank, C., et al. (2018). A Revised Diagnostic Classification of Canine Glioma: Towards Validation of the Canine Glioma Patient as a Naturally Occurring Preclinical Model for Human Glioma. J Neuropathol Exp Neurol 77, 1039–1054.

LeBlanc, A. K., Mazcko, C., Brown, D. E., Koehler, J. W., Miller, A. D., Miller, C. R., Bentley, R. T., Packer, R. A., Breen, M., Boudreau, C. E., et al. (2016). Creation of an NCI comparative brain tumor consortium: informing the translation of new knowledge from canine to human brain tumor patients. Neuro Oncol 18, 1209–1218.

Lindblad-Toh, K., Garber, M., Zuk, O., Lin, M. F., Parker, B. J., Washietl, S., Kheradpour, P., Ernst, J., Jordan, G., Mauceli, E., et al. (2011). A high-resolution map of human evolutionary constraint using 29 mammals. Nature 478, 476.

Louis, D. N., Perry, A., Reifenberger, G., von Deimling, A., Figarella-Branger, D., Cavenee, W. K., Ohgaki, H., Wiestler, O. D., Kleihues, P., and Ellison, D. W. (2016). The 2016 World Health Organization Classification of Tumors of the Central Nervous System: a summary. Acta Neuropathol 131, 803–820.

Ma, X., Liu, Y., Liu, Y., Alexandrov, L. B., Edmonson, M. N., Gawad, C., Zhou, X., Li, Y., Rusch, M. C., Easton, J., et al. (2018). Pan-cancer genome and transcriptome analyses of 1,699 paediatric leukaemias and solid tumours. Nature.

Mackay, A., Burford, A., Carvalho, D., Izquierdo, E., Fazal-Salom, J., Taylor, K. R., Bjerke, L., Clarke, M., Vinci, M., Nandhabalan, M., et al. (2017). Integrated Molecular Meta-Analysis of 1,000 Pediatric High-Grade and Diffuse Intrinsic Pontine Glioma. Cancer Cell 0.

Mansour, T. A., Lucot, K., Konopelski, S. E., Dickinson, P. J., Sturges, B. K., Vernau, K. L., Choi, S., Stern, J. A., Thomasy, S. M., Doring, S., et al. (2018). Whole genome variant association across 100 dogs identifies a frame shift mutation in DISHEVELLED 2 which contributes to Robinow-like syndrome in Bulldogs and related screw tail dog breeds. PLoS Genet 14, e1007850.

Martincorena, I., Raine, K. M., Gerstung, M., Dawson, K. J., Haase, K., Van Loo, P., Davies, H., Stratton, M. R., and Campbell, P. J. (2017). Universal Patterns of Selection in Cancer and Somatic Tissues. Cell 171, 1029–1041 e1021.

Newman, A. M., Liu, C. L., Green, M. R., Gentles, A. J., Feng, W., Xu, Y., Hoang, C. D., Diehn, M., and Alizadeh, A. A. (2015). Robust enumeration of cell subsets from tissue expression profiles. Nat Methods 12, 453–457.

Nowell, P. C. (1976). The clonal evolution of tumor cell populations. Science 194, 23–28.

Pai, A. A., Bell, J. T., Marioni, J. C., Pritchard, J. K., and Gilad, Y. (2011). A genome-wide study of DNA methylation patterns and gene expression levels in multiple human and chimpanzee tissues. PLoS Genet 7, e1001316.

Parker, H. G., VonHoldt, B. M., Quignon, P., Margulies, E. H., Shao, S., Mosher, D. S., Spady, T. C., Elkahloun, A., Cargill, M., Jones, P. G., et al. (2009). An expressed fgf4 retrogene is associated with breed-defining chondrodysplasia in domestic dogs. Science 325, 995–998.

Rao, R. C., and Dou, Y. (2015). Hijacked in cancer: the KMT2 (MLL) family of methyltransferases. Nat Rev Cancer 15, 334–346.

Sakthikumar, S., Elvers, I., Kim, J., Arendt, M. L., Thomas, R., Turner-Maier, J., Swofford, R., Johnson, J., Schumacher, S. E., Alföldi, J., et al. (2018). SETD2 Is Recurrently Mutated in Whole-Exome Sequenced Canine Osteosarcoma. Cancer Res 78, 3421–3431.

Schneider, G., Schmidt-Supprian, M., Rad, R., and Saur, D. (2017). Tissue-specific tumorigenesis: context matters. Nat Rev Cancer 17, 239–253.

Shay, T., Jojic, V., Zuk, O., Rothamel, K., Puyraimond-Zemmour, D., Feng, T., Wakamatsu, E., Benoist, C., Koller, D., Regev, A., and ImmGen, C. (2013). Conservation and divergence in the transcriptional programs of the human and mouse immune systems. Proc Natl Acad Sci U S A 110, 2946–2951.

Shinde, J., Bayard, Q., Imbeaud, S., Hirsch, T. Z., Liu, F., Renault, V., Zucman-Rossi, J., and Letouze, E. (2018). Palimpsest: an R package for studying mutational and structural variant signatures along clonal evolution in cancer. Bioinformatics 34, 3380–3381.

Snyder, J. M., Shofer, F. S., Van Winkle, T. J., and Massicotte, C. (2006). Canine intracranial primary neoplasia: 173 cases (1986-2003). J Vet Intern Med 20, 669–675.

Song, R. B., Vite, C. H., Bradley, C. W., and Cross, J. R. (2013). Postmortem evaluation of 435 cases of intracranial neoplasia in dogs and relationship of neoplasm with breed, age, and body weight. J Vet Intern Med 27, 1143–1152.

Stearns, S. C. (1992). The evolution of life histories, (Oxford; New York: Oxford University Press).

Sturm, D., Bender, S., Jones, D. T., Lichter, P., Grill, J., Becher, O., Hawkins, C., Majewski, J., Jones, C., Costello, J. F., et al. (2014). Paediatric and adult glioblastoma: multiform (epi)genomic culprits emerge. Nat Rev Cancer 14, 92–107.

Targa, A., and Rancati, G. (2018). Cancer: a CINful evolution. Curr Opin Cell Biol 52, 136–144.

Tate, J. G., Bamford, S., Jubb, H. C., Sondka, Z., Beare, D. M., Bindal, N., Boutselakis, H., Cole, C. G., Creatore, C., Dawson, E., et al. (2019). COSMIC: the Catalogue Of Somatic Mutations In Cancer. Nucleic Acids Res 47, D941–D947.

Taylor, A. M., Shih, J., Ha, G., Gao, G. F., Zhang, X., Berger, A. C., Schumacher, S. E., Wang, C., Hu, H., Liu, J., et al. (2018). Genomic and Functional Approaches to Understanding Cancer Aneuploidy. Cancer Cell 33, 676–689 e673.

TCGA_Network, Brat, D. J., Verhaak, R. G., Aldape, K. D., Yung, W. K., Salama, S. R., Cooper, L. A., Rheinbay, E., Miller, C. R., Vitucci, M., et al. (2015). Comprehensive, Integrative Genomic Analysis of Diffuse Lower-Grade Gliomas. N Engl J Med 372, 2481–2498.

Thompson, M. J., vonHoldt, B., Horvath, S., and Pellegrini, M. (2017). An epigenetic aging clock for dogs and wolves. Aging (Albany NY) 9, 1055–1068.

Tollis, M., Schiffman, J. D., and Boddy, A. M. (2017). Evolution of cancer suppression as revealed by mammalian comparative genomics. Curr Opin Genet Dev 42, 40–47.

Truve, K., Dickinson, P., Xiong, A., York, D., Jayashankar, K., Pielberg, G., Koltookian, M., Muren, E., Fuxelius, H. H., Weishaupt, H., et al. (2016). Utilizing the Dog Genome in the Search for Novel Candidate Genes Involved in Glioma Development-Genome Wide Association Mapping followed by Targeted Massive Parallel Sequencing Identifies a Strongly Associated Locus. PLoS Genet 12, e1006000.

Varn, F. S., Wang, Y., Mullins, D. W., Fiering, S., and Cheng, C. (2017). Systematic Pan-Cancer Analysis Reveals Immune Cell Interactions in the Tumor Microenvironment. Cancer Res 77, 1271–1282.

Venkatesan, S., Birkbak, N. J., and Swanton, C. (2017). Constraints in cancer evolution. Biochem Soc Trans 45, 1–13.

Villar, D., Berthelot, C., Aldridge, S., Rayner, T. F., Lukk, M., Pignatelli, M., Park, T. J., Deaville, R., Erichsen, J. T., Jasinska, A. J., et al. (2015). Enhancer evolution across 20 mammalian species. Cell 160, 554–566.

Wong, K., van der Weyden, L., Schott, C. R., Foote, A., Constantino-Casas, F., Smith, S., Dobson, J. M., Murchison, E. P., Wu, H., Yeh, I., et al. (2019). Cross-species genomic landscape comparison of human mucosal melanoma with canine oral and equine melanoma. Nat Commun 10, 353.

## Supplemental references

Angermueller, C., Lee, H. J., Reik, W., and Stegle, O. (2017). DeepCpG: accurate prediction of single-cell DNA methylation states using deep learning. Genome Biology 18.

Barthel, F. P., Wei, W., Tang, M., Martinez-Ledesma, E., Hu, X., Amin, S. B., Akdemir, K. C., Seth, S., Song, X., Wang, Q., et al. (2017). Systematic analysis of telomere length and somatic alterations in 31 cancer types. Nat Genet 49, 349–357.

Broeckx, B. J., Hitte, C., Coopman, F., Verhoeven, G. E., De Keulenaer, S., De Meester, E., Derrien, T., Alfoldi, J., Lindblad-Toh, K., Bosmans, T., et al. (2015). Improved canine exome designs, featuring ncRNAs and increased coverage of protein coding genes. Sci Rep 5, 12810.

Chen, S., Zhou, Y., Chen, Y., and Gu, J. (2018). fastp: an ultra-fast all-in-one FASTQ preprocessor. Bioinformatics 34, i884–i890.

Cibulskis, K., Lawrence, M. S., Carter, S. L., Sivachenko, A., Jaffe, D., Sougnez, C., Gabriel, S., Meyerson, M., Lander, E. S., and Getz, G. (2013). Sensitive detection of somatic point mutations in impure and heterogeneous cancer samples. Nat Biotechnol 31, 213–219.

DePristo, M. A., Banks, E., Poplin, R., Garimella, K. V., Maguire, J. R., Hartl, C., Philippakis, A. A., del Angel, G., Rivas, M. A., Hanna, M., et al. (2011). A framework for variation discovery and genotyping using next-generation DNA sequencing data. Nat Genet 43, 491–498.

Deshwar, A. G., Vembu, S., Yung, C. K., Jang, G. H., Stein, L., and Morris, Q. (2015). PhyloWGS: reconstructing subclonal composition and evolution from whole-genome sequencing of tumors. Genome Biol 16, 35.

Fang, L. T., Afshar, P. T., Chhibber, A., Mohiyuddin, M., Fan, Y., Mu, J. C., Gibeling, G., Barr, S., Asadi, N. B., Gerstein, M. B., et al. (2015). An ensemble approach to accurately detect somatic mutations using SomaticSeq. Genome Biol 16, 197.

Farmery, J. H. R., Smith, M. L., Diseases, N. B.-R., and Lynch, A. G. (2018). Telomerecat: A ploidy-agnostic method for estimating telomere length from whole genome sequencing data. Sci Rep 8, 1300.

Fleshner, I., and Chernett, N. L. (1997). A wellness model for the geriatric population. Home Care Provid 2, 321–323.

Gentles, A. J., Newman, A. M., Liu, C. L., Bratman, S. V., Feng, W., Kim, D., Nair, V. S., Xu, Y., Khuong, A., Hoang, C. D., et al. (2015). The prognostic landscape of genes and infiltrating immune cells across human cancers. Nat Med 21, 938–945.

Gerstung, M., Jolly, C., Leshchiner, I., Dentro, S. C., Gonzalez, S., Mitchell, T. J., Rubanova, Y., Anur, P., Rosebrock, D., Yu, K., et al. (2017). The evolutionary history of 2,658 cancers. bioRxiv.

Ha, G., Roth, A., Khattra, J., Ho, J., Yap, D., Prentice, L. M., Melnyk, N., McPherson, A., Bashashati, A., Laks, E., et al. (2014). TITAN: inference of copy number architectures in clonal cell populations from tumor whole-genome sequence data. Genome Res 24, 1881–1893.

Imielinski, M., Berger, A. H., Hammerman, P. S., Hernandez, B., Pugh, T. J., Hodis, E., Cho, J., Suh, J., Capelletti, M., Sivachenko, A., et al. (2012). Mapping the hallmarks of lung adenocarcinoma with massively parallel sequencing. Cell 150, 1107–1120.

Iorio, F., Garcia-Alonso, L., Brammeld, J. S., Martincorena, I., Wille, D. R., McDermott, U., and Saez-Rodriguez, J. (2018). Pathway-based dissection of the genomic heterogeneity of cancer hallmarks’ acquisition with SLAPenrich. Sci Rep 8, 6713.

Koboldt, D. C., Larson, D. E., and Wilson, R. K. (2013). Using VarScan 2 for Germline Variant Calling and Somatic Mutation Detection. Curr Protoc Bioinformatics 44, 15 1411–17.

Koster, J., and Rahmann, S. (2018). Snakemake-a scalable bioinformatics workflow engine. Bioinformatics 34, 3600.

Krueger, F., and Andrews, S. R. (2011). Bismark: a flexible aligner and methylation caller for Bisulfite-Seq applications. Bioinformatics 27, 1571–1572.

Lee, S., Lee, S., Ouellette, S., Park, W. Y., Lee, E. A., and Park, P. J. (2017). NGSCheckMate: software for validating sample identity in next-generation sequencing studies within and across data types. Nucleic Acids Res 45, e103.

McGranahan, N., Favero, F., de Bruin, E. C., Birkbak, N. J., Szallasi, Z., and Swanton, C. (2015). Clonal status of actionable driver events and the timing of mutational processes in cancer evolution. Sci Transl Med 7, 283ra254.

McKenna, A., Hanna, M., Banks, E., Sivachenko, A., Cibulskis, K., Kernytsky, A., Garimella, K., Altshuler, D., Gabriel, S., Daly, M., and DePristo, M. A. (2010). The Genome Analysis Toolkit: a MapReduce framework for analyzing next-generation DNA sequencing data. Genome Res 20, 1297–1303.

McLaren, W., Gil, L., Hunt, S. E., Riat, H. S., Ritchie, G. R., Thormann, A., Flicek, P., and Cunningham, F. (2016). The Ensembl Variant Effect Predictor. Genome Biol 17, 122.

Mermel, C. H., Schumacher, S. E., Hill, B., Meyerson, M. L., Beroukhim, R., and Getz, G. (2011). GISTIC2.0 facilitates sensitive and confident localization of the targets of focal somatic copy-number alteration in human cancers. Genome Biol 12, R41.

Okonechnikov, K., Conesa, A., and Garcia-Alcalde, F. (2016). Qualimap 2: advanced multi-sample quality control for high-throughput sequencing data. Bioinformatics 32, 292–294.

Parker, M., Chen, X., Bahrami, A., Dalton, J., Rusch, M., Wu, G., Easton, J., Cheung, N. K., Dyer, M., Mardis, E. R., et al. (2012). Assessing telomeric DNA content in pediatric cancers using whole-genome sequencing data. Genome Biol 13, R113.

Pimentel, H., Bray, N. L., Puente, S., Melsted, P., and Pachter, L. (2017). Differential analysis of RNA-seq incorporating quantification uncertainty. Nat Methods 14, 687–690.

Venkatesan, S., and Swanton, C. (2016). Tumor Evolutionary Principles: How Intratumor Heterogeneity Influences Cancer Treatment and Outcome. Am Soc Clin Oncol Educ Book 35, e141–149.

Vladoiu, M. C., El-Hamamy, I., Donovan, L. K., Farooq, H., Holgado, B. L., Sundaravadanam, Y., Ramaswamy, V., Hendrikse, L. D., Kumar, S., Mack, S. C., et al. (2019). Childhood cerebellar tumours mirror conserved fetal transcriptional programs. Nature.

Wilm, A., Aw, P. P., Bertrand, D., Yeo, G. H., Ong, S. H., Wong, C. H., Khor, C. C., Petric, R., Hibberd, M. L., and Nagarajan, N. (2012). LoFreq: a sequence-quality aware, ultra-sensitive variant caller for uncovering cell-population heterogeneity from high-throughput sequencing datasets. Nucleic Acids Res 40, 11189–11201.

